# Leader cell activity and collective invasion by an autocrine nucleotide loop through connexin-43 hemichannels and ADORA1

**DOI:** 10.1101/2019.12.30.888958

**Authors:** Antoine A. Khalil, Olga Ilina, Angela Vasaturo, Jan-Hendrik Venhuizen, Manon Vullings, Victor Venhuizen, Ab Bilos, Carl Figdor, Paul N. Span, Peter Friedl

## Abstract

Progression of epithelial cancers predominantly proceeds by collective invasion of cell groups with coordinated cell-cell junctions and multicellular cytoskeletal activity. Collectively invading breast cancer cells co-express adherens junctions and connexin-43 (Cx43) gap junctions *in vitro* and in patient samples, yet whether gap junctions contribute to collective invasion remains unclear. We here show that Cx43 is required for chemical coupling between collectively invading breast cancer cells and, by its hemichannel function, adenosine nucleotide release into the extracellular space. Using molecular interference and rescue strategies *in vitro* and in orthotopic mammary tumors *in vivo*, Cx43-dependent adenosine nucleotide release was identified as essential mediator engaging the nucleoside receptor ADORA1, to induce leader cell activity and collective migration. In clinical samples joint-upregulation of Cx43 and ADORA1 predicts decreased relapse-free survival. This identifies autocrine nucleotide signaling, through a Cx43/ADORA1 axis, as critical pathway in leader cell function and collective cancer cell invasion.

**Graphical abstract:** 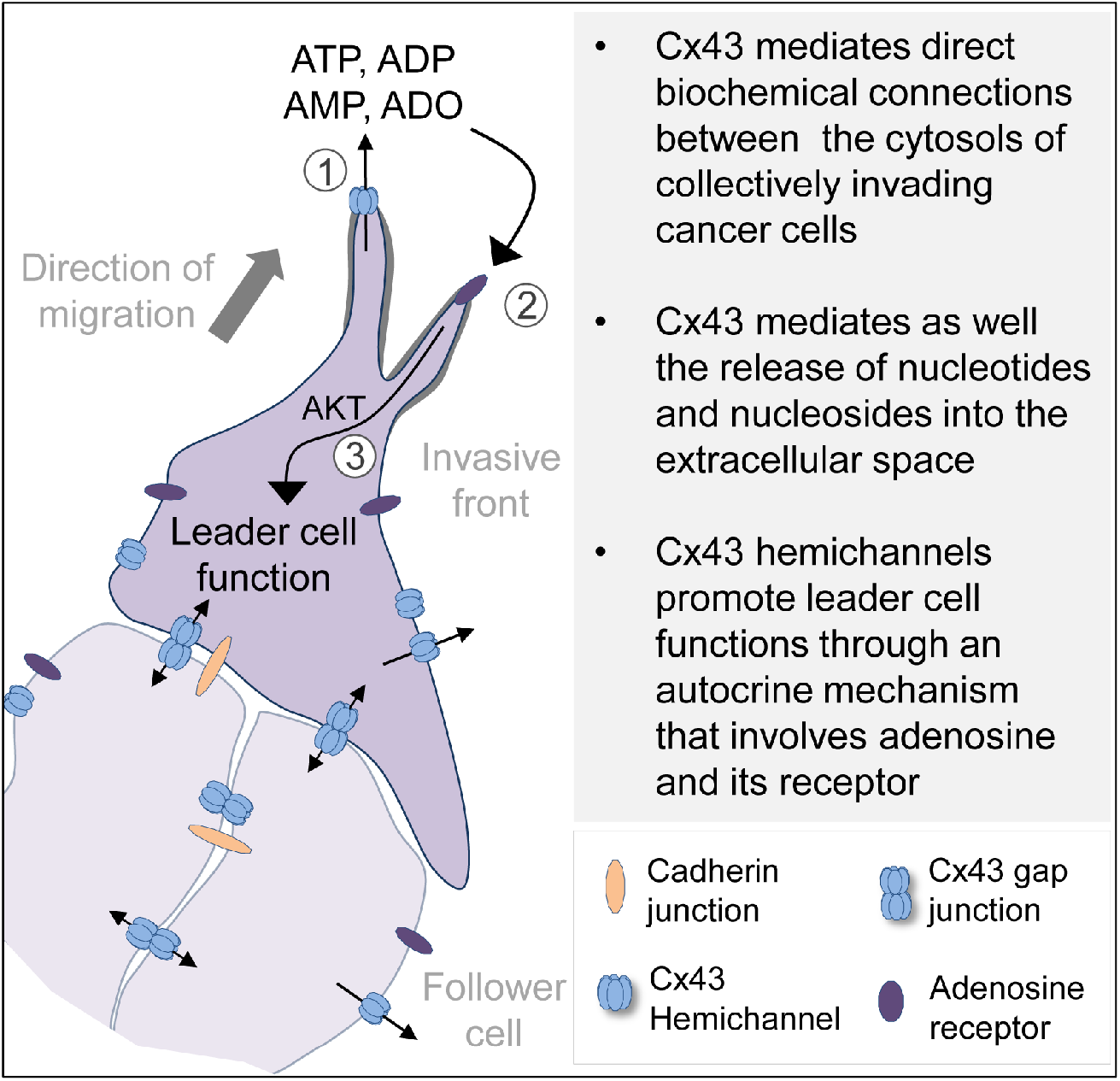

## Introduction

The collective invasion of multicellular groups with intact cell-cell junctions is an important mechanism to local invasion and metastasis in epithelial cancers (Cheung and Ewald, 2016; Gaggioli et al., 2007; Khalil et al., 2017). Collective invasion is initiated and maintained by leader cells that extend actin-rich protrusions, engage with the extracellular matrix (ECM) to exert traction and ECM remodeling, and maintain cell-cell coupling with follower cells (Cheung et al., 2013; Westcott et al., 2015). Mechanical connection and cell-to-cell signaling between moving cells are provided by adherens junctions, through E-, N- and other cadherins (Theveneau and Mayor, 2012). Cadherins coordinate the actomyosin cytoskeleton through catenins and vinculin (Hartsock and Nelson, 2008) and secure supracellular force balance, front-rear and apicobasal polarity, and multicellular branching (Tambe et al., 2011).

In non-neoplastic collective processes during morphogenesis and regeneration, moving cell sheets are further connected by gap junctions (Ashton et al., 1999; Huang et al., 1998; Kotini et al., 2018; Marins et al., 2009). Gap junctions consist of connexins (Cx) oligomerized into hemichannels that engage across cell membranes (Goodenough and Paul, 2009). The resulting transmembrane connections mediate intercellular transfer of ions and small molecules (<1kDa), including Ca^++^, phosphoinosites and nucleotides (Goodenough and Paul, 2009). Connexins mediate multicellular contractility of cardiomyocytes and the myoepithelial layer of mammary ducts (Kumai et al., 2000; Mroue et al., 2015) and, by unknown mechanisms, collective movement of neural and endothelial cells (Ashton et al., 1999; Huang et al., 1998; Marins et al., 2009). The mechanisms by which connexins support coordinated cytoskeletal contractility and multicellular dynamics vary. Direct cell-to-cell signaling occurs through gap-junctional intercellular transfer of second messengers, including IP3 and cAMP, energy equivalents (ATP, glucose), and Ca^2+^ wave propagation (Boitano et al., 1992; Goldberg et al., 2004; Howe, 2004). Connexins further contribute to gene expression of cadherins (Kotini et al., 2018) and/or the release of chemotactic factors to induce cell polarity and migration (Barletta et al., 2012; Haynes et al., 2006; Kaczmarek et al., 2005). In morphogenesis, connexins are indispensable for coordinated tissue growth (Sinyuk et al., 2018). By regulating cell differentiation, connexins further counteract neoplastic transformation (Bazzoun et al., 2019; Fostok et al., 2019; Saunders, 2001; Zhang et al., 2003) and forced expression of connexins in transformed cells reduces tumor cell growth and inhibits invasion by reverting the epithelial-to-mesenchymal transition (Kazan et al., 2019; McLachlan et al., 2006). On the other hand, connexins remain expressed in several solid tumors and expression increases in metastases (Bos et al., 2009; Elzarrad et al., 2008; Kanczuga-Koda et al., 2006; Stoletov et al., 2013). Interference with Cx43 expression or channel function inhibits cancer cell migration *in vitro* (Ogawa et al., 2012) and reduces metastatic seeding by reducing binding of circulating tumor cells to vascular endothelial cells (el-Sabban and Pauli, 1991; Elzarrad et al., 2008; Stoletov et al., 2013). Because connexins exert a range of functions, and these may vary between tumor types and experimental conditions, an integrating concept on how connexins either suppress or enhance neoplastic progression is lacking (Aasen et al., 2019; Naus and Laird, 2010).

Whereas a role of connexins in collective processes in morphogenesis has been established, the contribution of gap junctional communication in collective tumor cell invasion remains unclear (Friedl and Gilmour, 2009). We here revisited Cx43 expression and function in breast cancer models of collective invasion. Using pharmacological inhibition, molecular interference and rescue strategies, we show that connexins are required for collective polarity and invasion *in vitro* and *in vivo*. Cx43-mediated communication between collectively invading cancer cells and connexin hemichannels are required to release adenosine nucleotides which initiates an autocrine loop inducing leader cell function and collective guidance through adenosine receptor 1 (ADORA1).

## Results

### Breast cancer cells express Cx43 during collective invasion

To address whether connexins are expressed in invasive ductal breast carcinoma infiltrating the fibrous tumor stroma and adipose tissue, Cx43 expression was detected in patient samples by multispectral imaging (Suppl. Fig. 1a, b) (Mascaux et al., 2019). Non-neoplastic ducts consisted of luminal epithelial cells with negative to low Cx43 expression and Cx43-positive myoepithelial cells (Fig. 1a, b; Suppl. Fig 1c). In tumors, however, multicellular epithelial strands or roundish clusters with cytokeratin expression located in the fibrous or adipose tissue showed increased Cx43 expression in 9/12 samples, and expression was increased compared to normal luminal epithelium (Fig. 1a, arrowheads; Fig. 1c; Suppl. Fig. 1c, d). The fraction of Cx43^bright^ cells, as quantified by single-cell *in situ* cytometry in Cx43-positive samples (Fig. 1c, red label), varied from 5-75% (Suppl. Fig. 1d). This indicates that invasive ductal carcinomas contain subregions of high Cx43 expression in the invasion zones. Using meta-analysis of the Gyorffy cohort of basal-type breast cancer patients (Gyorffy et al., 2010), high Cx43 gene expression was associated with reduced relapse-free survival (Fig. 1d), but not distant metastasis-free survival (Suppl. Fig. 1e).

**Figure 1.**
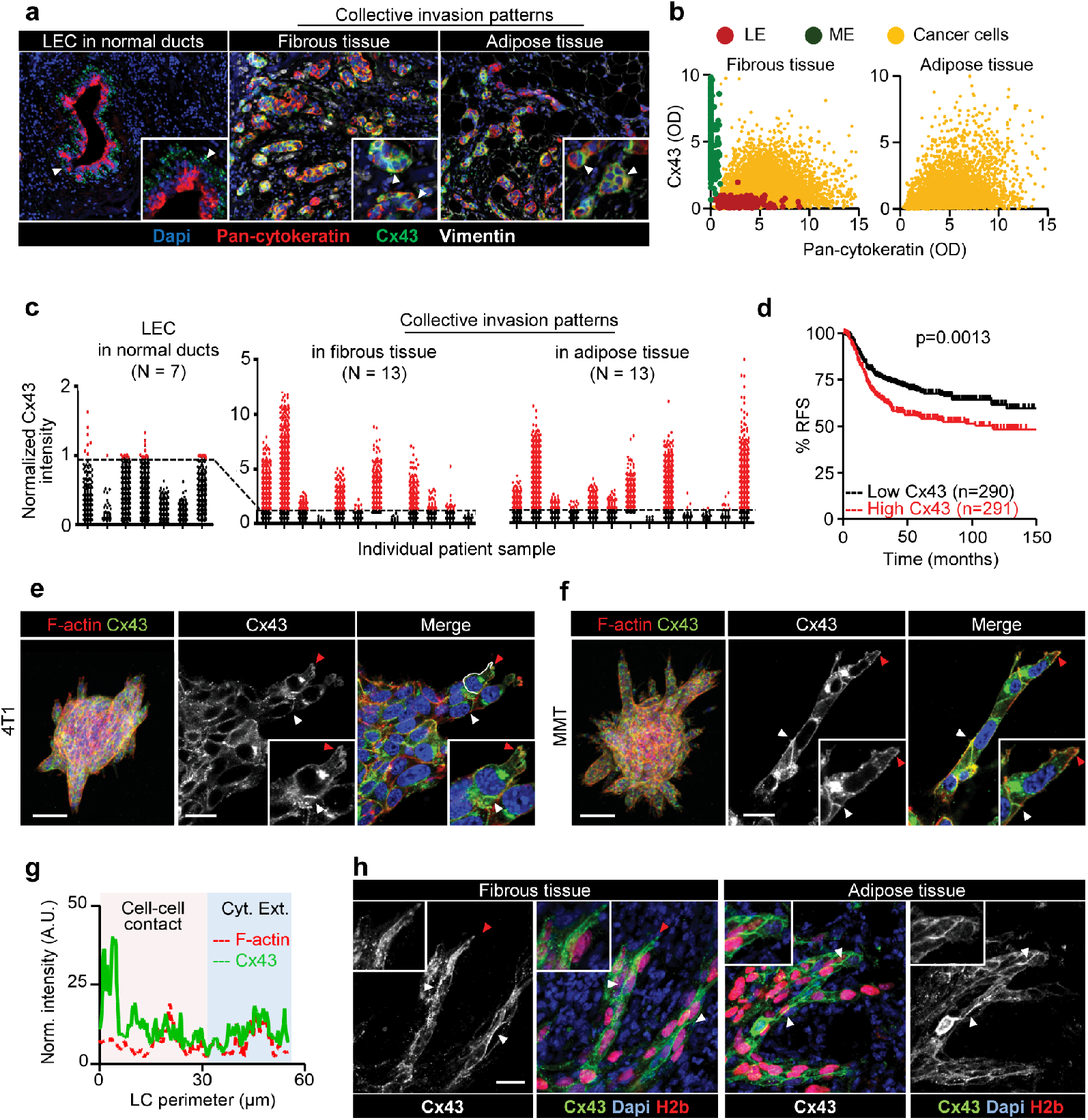
Increased expression of Cx43 in breast cancer cells during collective cancer cell invasion in vivo and in vitro. (a) Multispectral microscopy of breast cancer tissue sections stained for Cx43 (green), pan cytokeratin (red; epithelial cells), vimentin (white; stromal cells, cancer cells after EMT). The three panels represent example slices for normal ducts and the invasion zones in the collagen-rich fibrous or adipose tissue. Detection of Cx43 in myoepithelial cells (ME) surrounding the luminal epithelium (LE) and epithelial cancer cells along cell-cell junctions. White arrowheads, junctional Cx43. (b) Colocalization analysis of Cx43 vs pan cytokeratin expression and classified myoepithelial cells, luminal epithelial cells and invading cancer cells in fibrous or adipose tissue. Cx43 and pan cytokeratin levels were acquired using supervised automated tissue segmentation software (Inform). (c) Cx43 levels in LE cells, cancer cells within the fibrous or adipose tissue from 13 clinical samples (see Table 2 for sample details). Dotted line represents the threshold level for Cx43 negativity, determined by receiver operating characteristics (ROC) analysis with intensity values obtained from the luminal epithelial cells as control. (d) Kaplan-Meier survival plot predicting relapse free survival for high vs low Cx43 expression in basal-type breast cancer patients (Gyorffy B et al.). P values, Log-rank test. (e-f) Distribution of Cx43 along cell-cell junctions (white arrowheads) and at the polar extensions (red arrowheads) of leader cells during collective invasion of 4T1 and MMT spheroids in 3D collagen. (g) Intensity distribution of Cx43 and F-actin along the circumference of a leader cell (g, white dashed border). Values show the pixel intensity with background subtraction. Confocal images are displayed as maximum intensity projections from a 3D confocal stack. (h) Cx43 expression and distribution during collective invasion of 4T1 cells in the fibrous and adipose tissue of the mouse mammary gland monitored by confocal microscopy. Bars, 100 μm (g, overview), 20 μm (e, f, h).

To address whether Cx43 supports invasion, the metastatic breast cancer models 4T1, which expresses E-cadherin, and MMT, which lacks E-cadherin but expresses N-cadherin (Ilina et al., 2018), were embedded as multicellular spheroids in 3D collagen lattices and monitored for invasion over 24-48 h (Suppl. Movies 1, 2). 4T1 and MMT cells invaded as collective strands and expressed high levels of Cx43 and Cx45 mRNA and low levels of other connexins after isolation from collagen matrix culture (Suppl. Fig. 1f). During collective invasion, Cx43 was localized along cell-cell junctions (Fig. 1e, f, white arrowheads) and in non-junctional domains, along cell protrusions (Fig. 1e, f, red arrowheads; g). Junctional and non-junctional Cx43 localization was further observed during invasion into the mammary fat pad *in vivo* (Fig. 1h). In contrast to Cx43, Cx45 was absent along cell-cell junctions, but localized in the cytosol (Suppl. Fig. 1g). Thus, Cx43 is expressed by collectively invading cells *in vitro* and *in vivo*.

### GJIC communication during collective invasion

Collective invasion in 3D spheroid culture was initiated by single or few leader cells, which extended actin-rich protrusions to the front and maintained cadherin-based junctions with following cells (Suppl. Fig. 2a, b). To examine whether gap junctions are active during collective invasion, intercellular dye transfer was measured in 3D spheroid invasion cultures using gap fluorescence recovery after photobleaching (3D-gapFRAP) (Suppl. Movie 3; Fig. 2a). Within minutes after photobleaching, leader and follower cells recovered the calcein signal by 20-30% (Fig. 2b, c), with unbleached neighboring cells acting as donors (Suppl. Movie 3; Suppl. Fig. 2c). As internal control, detached single cells lacked dye transfer after photobleaching (Fig. 2b, c; Suppl. Fig. 2c, d, purple cell outline; Suppl. Movie 3). Fluorescence recovery was reduced by 70-85% after inhibition of connexin channel function by carbenoxolone (CBX) (Fig. 2b-d; Suppl. Movie 3). Fluorescence recovery was further impaired when Cx43 expression was stably downregulated (by >80%) using shRNA (Fig. 2e, f; Suppl. Fig 2e, f, g).

**Figure 2.**
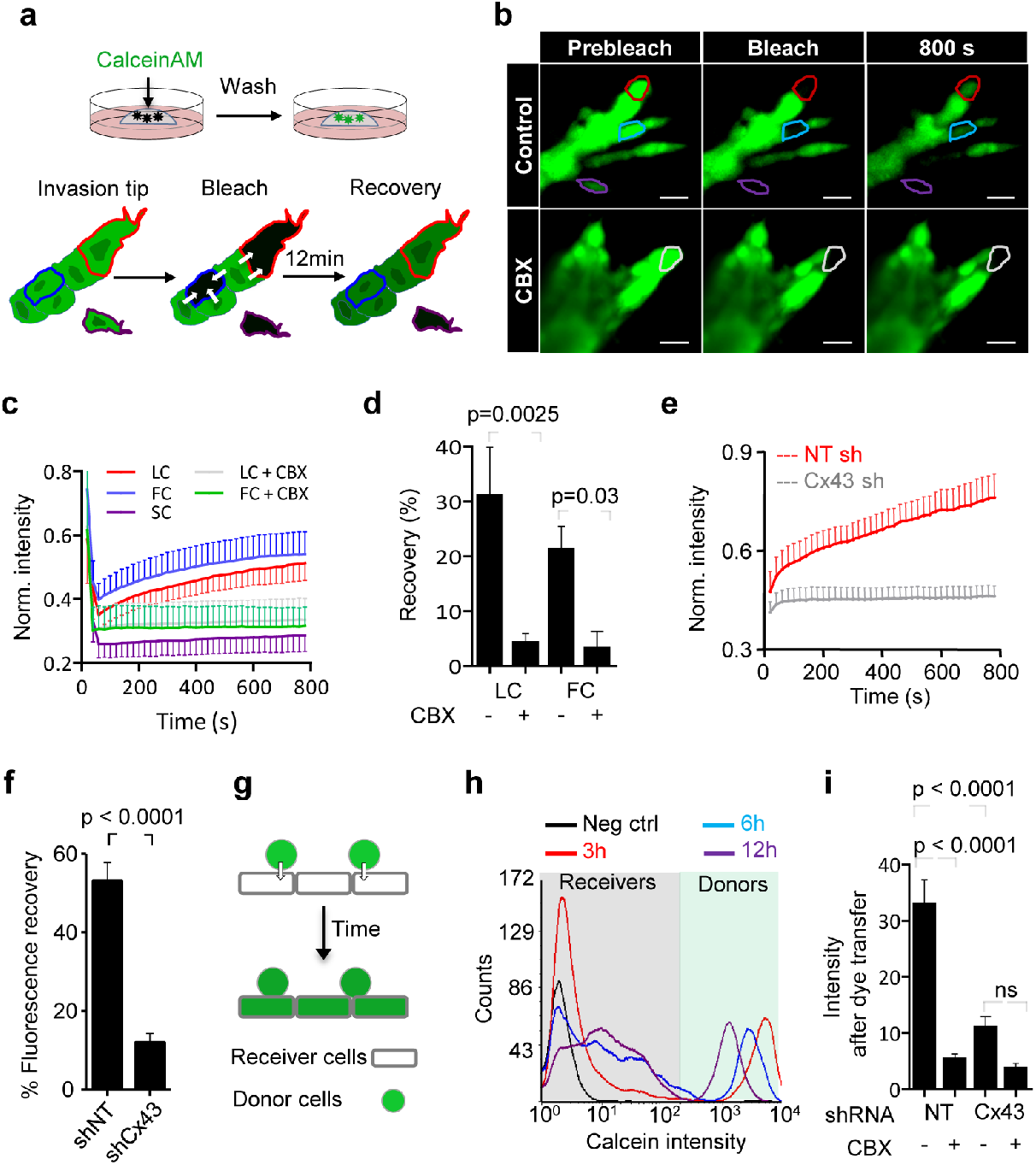
Collectively invading cancer cells maintain direct cell-cell communication through Cx43-mediated GJIC. (a) Workflow of time-resolved 3D gapFRAP, including (i) calcein labeling of multicellular spheroids in 3D collagen cultures and (ii) 3D gapFRAP procedure of leader and follower cells within a 3D invasion strand by asymmetric ROIs selection, photo-bleaching, followed by recording the change in fluorescence intensity over time. (b) Single confocal slices of calcein-labeled invasive 4T1 strands before and after photobleaching. Dashed contours represent the bleached leader cell (LC, red), follower cell (FC, blue), single detached cell (SC, purple) in control media and LC in the presence of CBX (white). (c) Normalized calcein fluorescence intensity in bleached cells in the presence or absence of CBX. Values represent normalized mean fluorescence intensities with SEM, 11-16 cells per treatment condition from 4 independent experiments (d) Effect of CBX on % fluorescence recovery after photobleaching of LC and FC. Values are represented as the means and SEM of 3 independent experiments. P values, Mann Whitney test. (e) Average fluorescence recovery in cells stably expressing control vector (shNT) or Cx43 shRNA (shCx43), during invasion in 3D collagen. Values represent the normalized mean intensities and SEM of 4 cells for each condition. (f) Inhibition of fluorescence recovery of MMT cells after Cx43 downregulation (cell groups in petri dish culture). Normalized mean intensity and SEM of 19-22 cells from 3 independent experiments. (g) Parachute assay to measure de novo junction formation followed by dye transfer into non-labeled cells (h) and example histograms obtained by flow cytometry showing the time-dependent alterations of calcein label in donor and recipient cells. (i) Mean intensity change of calcein uptake in receiver cells after 12 h of incubation with donor cells; in control conditions and during inhibition with CBX or after Cx43 down regulation. Mean intensities and SEM of 3 independent experiments. P values (f, i), Mann Whitney test. Bars, 20 μm.

To test whether all cells maintain GJIC, or only specific cell subsets, dye transfer between freshly interacting breast cancer cells was measured (parachute assay (Abbaci et al., 2008)) (Fig. 2g). Within a 12 h observation period, the majority of control cells (90-95%) received dye from the calcein-loaded monolayer (Fig. 2h). The extent of dye transfer was strongly reduced by 67-84% by CBX treatment or Cx43 downregulation (Fig. 2h; Suppl. Fig. 2h). This data indicates that collectively invading breast cancer cells interact by gap-junctional intercellular communication (GJIC) through Cx43.

### Cx43 is required for leader cell function

To test whether GJIC is required for collective cell invasion, 3D spheroid cultures were treated with pharmacological channel blockers CBX or 18 alpha-glycyrrhetinic acid (18aGA), at doses which did not compromise growth (Suppl. Fig. 3a, b). Cultures were treated either prior to (prevention) or after collective invasion has started (intervention). Both compounds dose-dependently reduced strand initiation and impaired the emergence of leader cells, compared to the inactive homologue GLZ or vehicle control (Fig. 3a, arrowheads, b, c; Suppl. Fig. 3c, d, arrowheads, e). Consequently, the invasion speed and cumulative length of collective invasion strands after 24 h were reduced 2-to 3-fold (Fig. 3b; Suppl. Fig. 3f-h).

**Figure 3.**
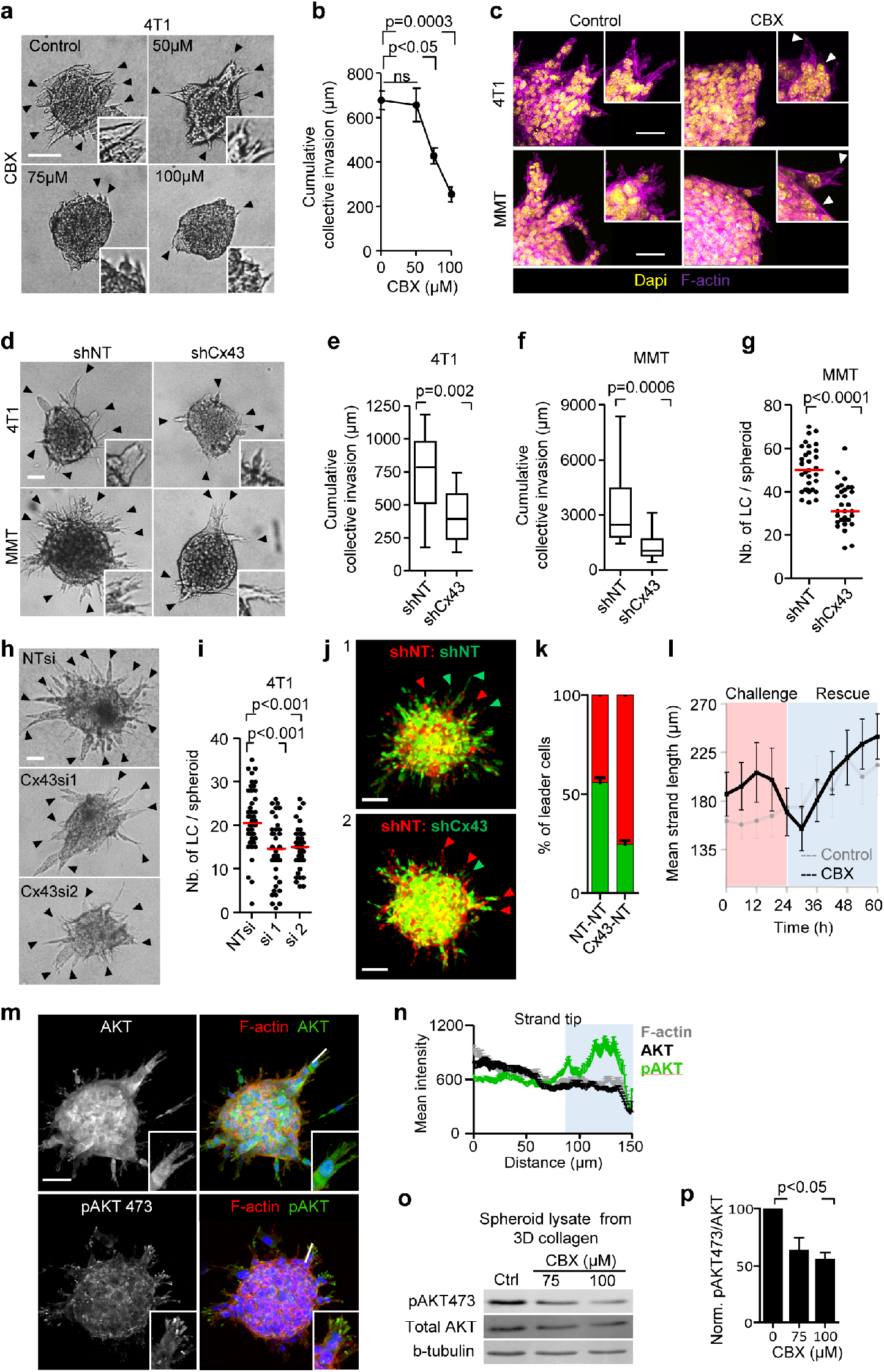
Cx43 dependence of leader cell function. (a) Bright field images of 4T1 spheroids after 24h of invasion in 3D collagen in the presence of increasing concentration of CBX. (b) Mean cumulative length of collective invasion strands in response to CBX. Mean values and SEM from 3 independent experiments each comprising 7 spheroids/condition. P values, ANOVA with Bonferoni multiple comparison test. (c) Maximum intensity projection from a confocal 3D stack, showing F-actin and DAPI in control and CBX treated spheroids (100μM). Bright field images (d), cumulative collective invasion (e, f) and leader cell initiation (g) of spheroids composed of 4T1 or MMT cells stably expressing Cx43 or non-targeting shRNA (NT). Values represent the medians from 3-4 independent experiments with 3-8 spheroids each. P values, Mann Whitney test. (h) Bright field images of 4T1 spheroids in response to transient Cx43 downregulation using two different Cx43 RNAi probes and (i) number of LCs per spheroid. (i) P values, ANOVA with Bonferoni multiple comparison test. (j) Maximum intensity projection from a confocal 3D stack showing mosaic spheroids composed of fluorescence-coded MMT cells stably expressing NT or Cx43 shRNA as indicated (1:1 ratio), arrowheads depict red or green leader cells. (k) Frequency of green or red leader cells. Bars represent the mean values and SEM pooled from 1 (j1) and 3 (j2) independent experiments. (l) Effect of CBX on the kinetics of established invasion strands. Mean length per invasion strand after addition of CBX at 0h (challenge) and wash out at 24 h (rescue). Data show the mean values and SEM from 6-8 invasion strands from one spheroid per condition. (m) Maximum intensity projection from a confocal 3D stack of MMT spheroids stained for AKT, phospho-AKT473 and F-actin. (n) Distribution of AKT and phospho-AKT473 and F-actin along extending protrusions of LCs (white dashed line in (m)). Line graphs represent the mean intensities of F-actin, AKT, and phospho-AKT473 from at least 12 leader cells. (o) Western blot and normalized intensities (p) for phospho-AKT473, AKT and β-tubulin as loading control. Whole cell lysates extracted from MMT spheroids after 24 h of invasion into 3D collagen in the presence of 0, 75 and 100 μM of CBX. Values represent normalized intensities of phospho-AKT473/total AKT with SD from 2 (75 μM) or 3 independent blots (0, 100 μM). P value, paired t-test. Bars, 50 μm (a, c, d, h), 25 μm (j, m).

Likewise, stable downregulation of Cx43 by shRNA compromised both the number and length of collectively invading strands (Fig. 3d-g). To rule out adaptation or off-target effects of stable Cx43 downregulation, transient Cx43 downregulation using individual RNAi probes was used, leading to reduced invasive strand initiation and elongation (Fig. 3h, i). Neither pharmacological nor RNA interference with GJIC altered cell-cell cohesion or shape. Invasion strands emerging despite interference retained intact morphology and cortical F-actin organization (Fig. 3c, arrowheads; Fig. 3d).

The decreased initiation and prolongation of invasion strands after interference with Cx43 suggested that the activity of leader cells was compromised. Leader cells are critical in initiating and maintaining collective invasion (Cheung et al., 2013; Zhang et al., 2019). To test whether leader cell function depends on Cx43, fluorescent mosaic spheroids of cells expressing non-targeting or Cx43 shRNA at equal ratio were allowed to invade. Cells with downregulated Cx43 were by 50% less likely to acquire leader cell position (Fig. 3j, k). CBX treatment reduced the speed of leader cells as assessed by cell tracking (Suppl. Fig. 3f; Suppl. Movie 4). Besides decreasing leader cell initiation, intermittent inhibition of GJIC by CBX in spheroids with established collective invasion affected leader cell function. CBX caused near-instantaneously collapsing protrusive tips, followed by speed reduction, and, ultimately, strand retraction (Fig. 3l; Suppl. Movie 5). Leader cell functions were restored after CBX washout (Suppl. Movie 5), indicating that connexins were required for the initiation and maintenance of collective invasion. In sheet migration of nontransformed epithelial cells, leader cell function depends on PI3K and AKT signaling (Yamaguchi et al., 2015). In 4T1 cells, the active form of AKT (AKT/pSer473) was abundant near the tips of leader-, but not in follower cells (Fig. 3m, n). Interference with GJIC by CBX decreased pAKT levels in invasion cultures by >40% (Fig. 3o, p), indicating that connexin channels are involved in AKT activation in leader cells.

### Cx43 hemichannels release purine nucleotides

In addition to its localization at cell-cell contacts, Cx43-positive foci were present in cell protrusions of leader cells (Fig. 1e, f, h, red arrowheads, g). Besides gap junctions, connexins can form transmembrane hemichannels (Ye, 2003), which release small molecules from the cytosol into the extracellular space, including glutamate, prostaglandins and the purine nucleotides ATP and ADP (Baroja-Mazo et al., 2013; Eltzschig et al., 2006; Retamal et al., 2007). Extracellular ATP and ADP and their degradation product adenosine (ADO), are important energy equivalents. In addition, extracellular nucleotides/nucleosides are effective extracellular signaling molecules, which induce cell polarization and migration in endothelial cells (Kaczmarek et al., 2005), microglia (Haynes et al., 2006), neutrophils (Barletta et al., 2012), as well as in individually moving cancer cells (Zhou et al., 2015). To address whether breast cancer cells release purine nucleotides during spheroid invasion culture, supernatants from 3D invasion cultures were analyzed for ADO, AMP, ADP and ATP by HPLC. Whereas cell-free media contained no purine metabolites, 3D spheroid invasion cultures after 24 h contained high levels of extracellular ATP (0.1 – 0.25 μg/mL) and moderate levels of ADO, AMP and ADP (10 – 50 ng/mL) (Fig. 4a). The concentration range reached *in vitro* was similar to ADO amounts in tumors *in vivo* (50-500 ng/ml) (de Andrade Mello et al., 2017). Inhibition of connexin channel-function by CBX and Cx43 knockdown both reduced the extracellular levels of ADO (by 26-42%), AMP (by 40-41%), ADP (by 60-64%) and ATP (by 28-52%) (Fig. 4b, c; Suppl. Fig 4a). This indicates that breast cancer cells release purine nucleosides and nucleotides through Cx43 hemichannels during 3D invasion culture.

**Figure 4:**
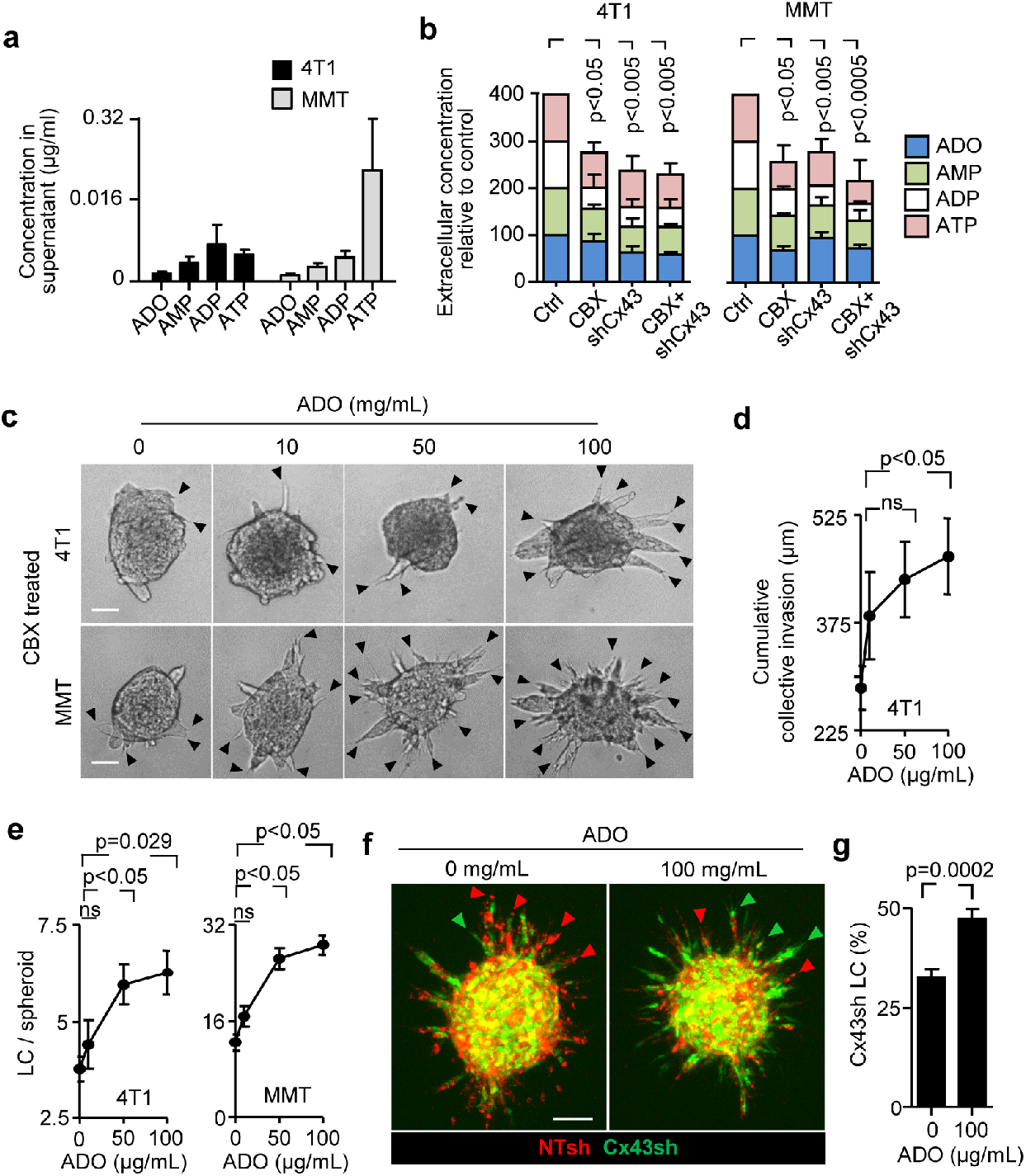
Cx43 hemichannel function in LC - purine nucleotides release. (a, b) HPLC analysis of purines released from 4T1 and MMT spheroids into the supernatant after 24h of invasion in 3D collagen. Values represent average concentrations of each purines (a) and the relative change of the total purine concentrations after treatment with CBX and or Cx43 downregulation in 4T1 and MMT cells (b). Mean values and SEM from 4 (a) or SD from 3 (b) independent experiments. P values, ANOVA Dunnett’s multiple comparison test. (c-e) Rescue of invasion in 4T1 and MMT spheroids in 3D collagen in the presence of CBX and escalating doses of ADO; for comparison with untreated baseline levels refer to Fig. 3b and Suppl. Fig. 3c, e. Representative bright-field images (c), cumulative collective invasion (d) and number of LC/spheroid (e). Mean values and SEM from 4 independent experiments each comprising of 5 to 11 spheroids/condition. P values, ANOVA with Bonferroni multiple comparison test. (f, g) Rescue of leader cell deficiency after Cx43 downregulation by exogenous ADO. (f) Maximum intensity projection from a 3D confocal stack showing mosaic spheroids with MMT cells stably expressing NT or Cx43 shRNA (1:1 ratio) in the absence or presence of ADO and (g) frequency of LC with green color expressing Cx43 shRNA. Bars represent the means and SEM from 3 independent experiments each comprising 3 to 6 spheroids. Bars, 50 μm (c), 100 μm (f).

In the extracellular space, ATP and ADP are converted to ADO, which in other models stimulates single-cell migration (Barletta et al., 2012; Stagg et al., 2010; Zhou et al., 2015). To test whether Cx43-dependent extracellular ADO also stimulates leader cell migration and, as a consequence, collective invasion of breast cancer cells, exogenous ADO was added to 4T1 and MMT spheroid cultures after leader cell function was compromised by CBX or Cx43 downregulation. In both 4T1 and MMT cells increasing concentrations of ADO partially (35-50%) restored the frequency of leader cells compared to untreated cells (Fig. 4c-e; Suppl. Fig. 4b). ADO substitution further restored defective leader cell function caused by downregulation of Cx43 in mosaic spheroids (Fig. 4f, green cells, g). The restored leader cell function by exogenous ADO thus indicates an autocrine Cx43 hemichannel-dependent purinergic signaling loop to secure collective migration.

### Autocrine purinergic receptor signaling maintains leader cell functions

The stimulating function of extracellular ADO in single-cell migration (Stagg et al., 2010; Zhou et al., 2015) depends on its binding to and signaling through P1 purinergic receptors (Peglion et al., 2014). P1 receptors belong to the G-protein coupled family of receptors, with the members ADORA1, ADORA2a, ADORA2b, ADORA3 (Antonioli et al., 2013). 4T1 and MMT cells isolated from 3D invasion culture expressed ADORA1 and ADORA2b mRNA but no other ADO receptor (Suppl. Fig. 4c). We thus tested whether ADORA1 or ADORA2b mediate collective invasion, using pharmacological interference. PSB36 and DPCPX, both selective ADORA1 antagonists with established activity *in vitro* and *in vivo* (Abo-Salem et al., 2004; Topfer et al., 2008), caused dose-dependent inhibition of collective invasion in 4T1 cells (Fig. 5a-c; Suppl. Fig. 4d) and MMT cells (Suppl. Fig. 4e). ADORA2b antagonist PSB1115 showed no effect on the emergence of leader cells of 4T1 and MMT cells (Suppl. Fig. 4f). Consistent with an impact on leader cell function, ADORA1 inhibition by PSB36 reduced AKT phosphorylation (Suppl. Fig. 4g), but did not affect the mitotic frequency of the cells (Suppl. Fig 4h). PSB36 abolished the effects of exogenous ADO on rescuing collective invasion (Fig 5d, e) and prevented leader cells to develop pointed anterior protrusions (Suppl. Movie 6). As a consequence of leader cell collapse, invasion strands retracted, and this was reversible after washout of the inhibitor (Suppl. Movie 6). Thus, leader cell function in collective invasion critically depends on ADO signaling through ADORA1.

**Figure 5:**
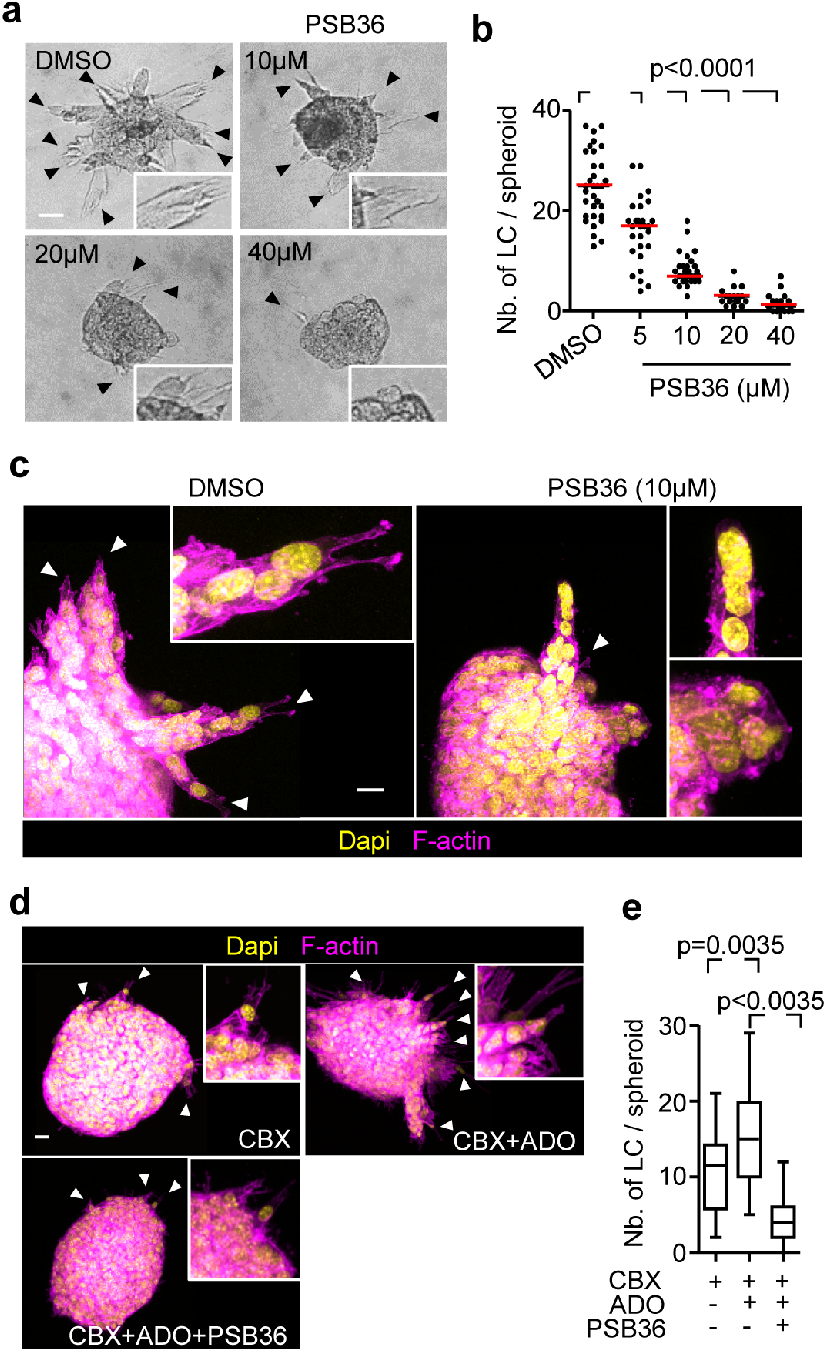
Autocrine purinergic receptor signaling maintains LC functions. (a) Brightfield images of 4T1 spheroids in 3D collagen for 24 h in the presence of escalating concentrations of PSB36. (b) Median numbers of LCs per spheroid (red line) from 3 independent experiments. P values, ANOVA with Bonferoni multiple comparison test. (c) Maximum intensity projection from a 3D confocal stack of 4T1 spheroid in the presence or absence of PSB36. White arrowheads, leading extensions of individual LCs. (d) Maximum intensity projections of 3D MMT spheroids with different treatments and (e) resulting number of leader cells per spheroid. Data represent the medians (black line), 25/75 percentiles (boxes) and maximum/minimum values (whiskers) from 2 independent experiments each comprising 10 to 16 spheroids/condition. P values, ANOVA with Bonferoni multiple comparison test. Bars, 50 μm (a), 25 μm (c d).

### ADORA1 in breast cancer invasion and progression *in vivo*

To address whether ADORA1 signaling supports collective breast cancer cell invasion *in vivo*, multicellular tumoroids were implanted into the mammary fat pad and the effect of systemic administration of PSB36 on progressing invasion was monitored by intravital multiphoton microscopy through a body window (Ilina et al., 2018). Untreated tumors invaded into the mammary fat pad via collective strands with elongated leading tip cells (Fig. 6a, arrowheads), as described (Ilina et al., 2018). PSB36 inhibited the extent of invasion with fewer strands with tips and reduced strand length (Fig. 6a, arrowheads; b). PSB36 did not affect tumor growth, assessed by the number of nuclei per spheroid (Fig. 6c), and this resulted in more compact tumors with increased packing density of cells. These data establish ADORA1 in promoting collective breast cancer cell invasion *in vivo*.

**Figure 6:**
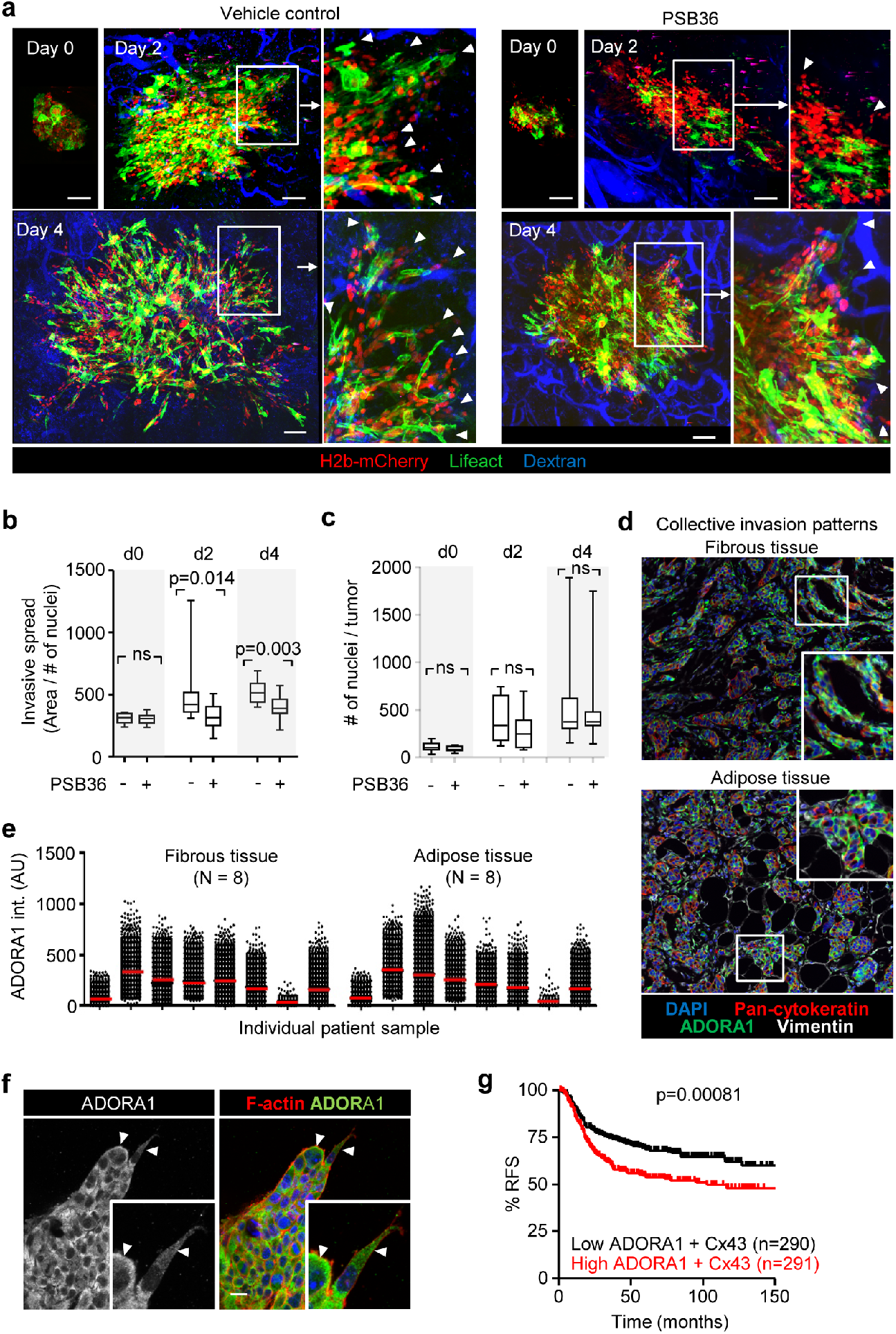
Pharmacological interference with ADORA1 inhibits collective invasion *in vivo*. (a) Invasion of 4T1 dual color cells (H2B/mCherry; LifeAct/YFP) from microtumors implanted into the mouse mammary fat pad and imaged by multiphoton microscopy through a mammary window at days 0, 2, and 4 post-implantation. Mice were treated with DMSO (vehicle) or 30mg/Kg of PSB36, intraperitoneal injection. (b, c) Analysis of invasion as the average area covered by the lesion normalized to the number of nuclei (b) and the tumor growth expressed as number of cells per lesion (c). Data in (b, c) represent the medians (black line), 25/75 percentiles (boxes) and maximum/minimum values (whiskers) from at least 9 implanted tumors from 4 independent experiments. P values, two-tailed unpaired Mann Whitney test. (d) Detection of ADORA1 in fibrous or adipose tissue invasion zones of breast cancer samples using multispectral microscopy. (e) Relative ADORA1 levels in epithelial cancer cells quantified from 8 independent lesions (see Table 2 for sample characteristics). (f) Distribution of ADORA1 in leader cells. Maximum intensity projection from 3D spheroid of 4T1 cells. Insets show the leading edge of strands. (g) Kaplan-Meier survival plot predicting relapse free survival (RFS) in basal-type breast cancer patients (Gyorffy B et al.) for high vs low ADORA1 expression combined with Cx43 expression. P values, Log-rank test. Bars, 100 μm (a), 25 μm (g).

In tumor lesions *in vivo*, extracellular ADO levels are elevated to micromolar ranges (Blay, 1997; Kumar, 2013) and ADORA1 mRNA and protein expression were reported to be increased in whole-tissue lysates of primary tumors compared to normal tissue (Khoo et al., 1996; Mirza et al., 2005). To verify that ADORA1 was also present in the invasion zone of ductal breast cancer, clinical samples were stained for ADORA1 and the collective invasion patterns located in the fibrous and adipose tissues were analyzed by multispectral microscopy (Fig. 6d). Multicellular strands and nests containing cytokeratin-positive epithelial cancer cells expressed ADORA1 (in 6/8 samples) with mostly cytosolic location (Fig. 6d, e). This distribution is consistent with predominantly cytosolic distribution of ADORA1 during collective invasion *in vitro* (Fig. 6f) and the reported cytoplasmic localization of ADORA1 and other chemokine receptors in nerve terminals and gastric cancer tissues (Iwasa et al., 2009; Rebola et al., 2003).

We finally tested whether inhibition of ADORA1 also reduces spontaneous metastasis to the lungs. 4T1 cells were implanted into the mammary fat pad and the mice treated daily with DMSO or PSB36. PSB36 therapy had no effect on the number or size of metastases to the lungs 30-36 days after tumor implantation (Suppl. Fig 5a-c). Thus, although PSB36 inhibited local invasion *in vivo*, it had no effect on spontaneous metastasis in the 4T1 model. ADORA1 expression was inversely correlated with relapse-free survival in the subset of patients with high Cx43 expression (Fig. 6g), but not ADORA1 alone (Suppl. Fig. 4i). This establishes the Cx43/ADORA1 axis as a prognostic parameter set for breast cancer progression.

## Discussion

We here identify an autocrine mechanism of leader cell activation and maintenance through Cx43 hemichannels, which release nucleotides to engage ADORA1 and induce collective invasion of breast cancer cells. ADORA1 drives local invasion and dissemination from implanted tumors in mice, and patients with high Cx43/ADORA1 expression suffer from worsened prognosis. Autocrine nucleotide signaling thus provides a cell-autonomous mechanism sustaining collective invasion in breast cancer and may deliver a rational basis for targeted intervention using hemichannel and ADORA inhibitors.

We here identified a previously unappreciated role of ADO in inducing leader cells that guide collective invasion of breast cancer cells, by cell polarization, adhesive interaction with ECM and ECM remodeling (Wolf et al., 2007). The nucleotide/ADORA signaling loops are known to regulate single-cell migration of normal and neoplastic cells (Chen et al., 2006; Ohsawa et al., 2012; Zhou et al., 2015). In the extracellular space, ATP and ADP are degraded into ADO by ectonucleotidases, including CD39 and CD73 (de Andrade Mello et al., 2017), and ADO levels in the high μM range are present in the tumor microenvironment (Blay et al., 1997); therefore ADO released in 3D invasion cultures and *in vivo* acts as both, metabolite and signaling effector of cancer progression (Antonioli et al., 2013). Leader cell induction by auto- or paracrine nucleotides depends on downstream signaling function through ADORAs to induce polarized cell-substrate interaction, and this is shared with nucleotide activity inducing single-cell migration in other contexts. AKT signaling, which is induced by G-protein coupled receptors (ADORA1, 2a, 2b, 3) mediates polarization of moving cells downstream of ADORA signaling (Chen et al., 2006; Gao et al., 2001; Othman et al., 2003; Stagg et al., 2010; Umapathy et al., 2013; Wen et al., 2011). Active AKT was increased in leader, but not follower cells and required both connexin channel and ADORA1 function. In single cell migration, individualized keratinocyte migration along electrical gradient depends on connexin/G-protein-coupled receptor axis (Riding and Pullar, 2015). In brain damage responses, astrocyte processes protrude in a connexin-ADORA-dependent manner to interact and close the microdefect (Davalos et al., 2005; Riding and Pullar, 2015). Although both ADORA1 and 2b are expressed, only ADORA1 mediated ADO-dependent invasion of 4T1 and MMT cells. Possibly, the reported 70fold higher affinity of ADO to ADORA1 (Fredholm et al., 2001) accounts for its dominance in leader-cell activation. However, in difference to ADORA-stimulated single-cell migration, autocrine stimulation of tip cells favors their engagement towards ECM while remaining coupled to follower cells, thus mediating the maintenance and elongation of collective invasion strands. The release of nucleotides by connexin hemichannels reveals a tumor-cell autonomous, autocrine mechanism of induction and continuation of collective invasion. The concept of hemichannel-mediated nucleotide release in our study is supported by: (i) focal Cx43 localization at cytoplasmic extensions of leader cells and lateral sites of follower cells; (ii) dependence of extracellular nucleotide levels in 3D tumor spheroid culture on Cx43 channel function; iii) the decreased probability of cells with low Cx43 expression to reach leader cell position; and (iv) the rapid loss of leader cell activity after connexin-channel inhibition or antagonization of ADORA1. These data strongly suggest Cx43 hemichannel-mediated nucleotide release as major pathway to collective invasion.

While ADORA1 function was clearly involved in local tissue invasion, spontaneous metastasis to the lungs was not affected by pharmacological ADORA1 antagonization. The 4T1 *in vivo* model resulted in 100% penetrance of spontaneous micro- and macrometastases and the used dosing of PSB36 inhibited invasion in mammary fat pads, suggesting that the drug is efficient *in vivo*. The lack of inhibition of metastases may indicate that leader cell function maintained by ADORA1 is of lower relevance in secondary organ colonization and metastatic outgrowth. A similar disconnect between local invasion and metastatic ability in distant organs was recently shown after deletion of E-cadherin, which caused increased local invasion in breast cancer models, but strongly compromised metastatic ability due to a survival defect (Padmanaban et al., 2019). Interfering with ADORA1 pathways may thus be justified to limit locally invasive disease, which cannot be effectively treated by surgery (Bakst et al., 2019). Beyond enhancing invasion, hemichannels and ADORAs also contribute to tumor growth and immunomodulation (Schalper et al., 2014), and orchestrate an integrated program to enhance local cancer progression. Purinergic signaling thus establishes a pro-invasive microenvironment through multipronged mechanisms, rendering pharmacological interference of nucleotide signaling through the connexin/ADORA axis in patients subsets as an attractive route to reprogram the tumor stroma and neoplastic invasion.

## Materials and Methods

### Antibodies and reagents

The following antibodies were used: rabbit anti-human Cx43 (3512, CST); chicken anti-human Vimentin (ab24525, Abcam); mouse anti-mouse β-Catenin (clone 14/ β -Catenin, BD Biosciences); rabbit anti-mouse AKT (9272, CST); rabbit anti-human phospho AKT Ser473 (4060, CST); mouse anti-human Pan Keratin (4545, CST); rabbit anti-rat ADORA1 (ab82477, Abcam); rabbit anti-human ADORA1 (ab124780, Abcam); Secondary polyclonal Alexa-fluor-488/647-conjugated goat anti-mouse, -rabbit or -chicken antibody. For F-actin visualization, Alexa-fluor-568-conjugated phalloidin (Invitrogen) was used. Adenosine (A4036, Sigma-Aldrich) was dissolved in water at a concentration of 1 mg/mL and stored in aliquots in - 20°C. The following inhibitors were used: gap junction channel inhibitors carbenoxolone disodium salt (CBX), 18-alpha glycyrrheitinic acid (18aGA) and the inactive inhibitor, glycyrrhizic acid ammonium salt (GLZ) (Sigma-Aldrich) (Kenny et al., 2002; Marins et al., 2009); ADORA1 selective and competitive antagonists 1-Butyl-3-(3-hydroxypropyl)-8-(3-noradamantyl) xanthine (PSB36) and 8-cyclopentyl-1,3-dipropylxanthine (DPCPX) (Sigma-Aldrich); ADORA2b selective antagonist 4-(2,3,6,7-Tetrahydro-2, 6-dioxo-1-propyl-1*H*-purin-8-yl)-benzenesulfonic acid (PSB1115) (Tocris) (Lin et al., 2010). CBX was dissolved in water in a stock concentration of 163 mM; all other inhibitors and antagonists were dissolved in DMSO in a stock concentration of 100-200 mM. All drugs were aliquoted into 5-20 μL and stored at - 20°C.

### Cell lines and culture

The following metastatic breast cancer cell lines were used: 4T1 (CRL-2539 ATCC); MMT wild type and MMT dual color cells (H2b-GFP and cytosolic RFP)(Tsuji et al., 2006) were a generous gift from R. Hoffman. Cells were maintained (37°C, 5% CO_2_ humidified atmosphere) in RPMI media (Invitrogen) supplemented with 10% fetal bovine serum (Sigma-Aldrich), penicillin (100 U/ml) and streptomycin (100 μg/ml; both PAA), L-glutamine (2 mM, Invitrogen). For generation of 4T1 and MMT cells with stable Cx43 RNA downregulation, mission lentiviral vector targeting mouse GJA1 (TRCN0000068473, sequence: CCGGCCCACCTTTGTGTCTTCCATACTCGAGTATGGAAGACACAAAGGTGGGTTTTTG) and non-targeting control, sequence (CCGGCAACAAGATGAAGAGCACCAACTCGAGTTGGTGCTCTTCATCTTGTTGTTTTT) was used (Sigma-Aldrich). For tumor implantation into the mammary fat pad *in vivo*, a stable 4T1/H2B-mCherry/LifeAct-GFP cells were used, as described (Ilina et al., 2018). Fluorescence expressing cells did not deviate from the wild-type counterparts in parameters, such as growth and invasion (data not shown).

### Cancer cell spheroid culture

Spheroids from 4T1 and MMT cells used in short-term *in vitro* migration assays were generated using the hanging drop method, as described (Ilina et al., 2018). Cells from subconfluent culture were detached with EDTA (1 mM) and trypsin (0.075%; Invitrogen), washed, suspended in medium/methylcellulose (2.4%; Sigma) and maintained as hanging droplets (25 μL) each containing 1000 cells. Hanging drops were kept in 37°C, 5% CO_2_ humidified atmosphere for 24 h (Del Duca et al., 2004). Spheroids were harvested and placed into DMEM media without FCS and without penicillin/streptomycin in a 6-well plate.

After cell aggregation, spheroids were washed in PBS and incorporated into 3D type I collagen lattices consisting of non-pepsinized rat-tail collagen (BD Biosciences) at a final concentration of 4 or 6 mg/ml for 4T1 or MMT cells, as described (Ilina et al., 2018). Only spheroids invading into the 3D phase of the collagen matrix were analyzed, whereas spheroids moving as 2D sheets underneath or on top of the culture were excluded.

### Transient downregulation of Cx43 by RNAi

4T1 and MMT cells were seeded in a 12-well plate at a density of 60×10^3^/well, in the presence of siRNA duplex (Dharmacon, Thermo Fisher Scientific). ON-Target plus non-targeting siRNA (D-001810-10) and two Cx43-specific targeting sequences (J-051694-05 and J-051694-06) were used at a final concentration of 50 nM using Dharmafect transfection reagent 4 (Thermo Scientific). After 12 h, RNAi duplex containing supernatant was removed and replaced by penicillin/streptomycin free media. After a recovery period of at least 6 h, cells were used for spheroid generation. For assessment of knockdown efficiency, whole-cell lysates were analyzed after 24, 48 and 72 h post-removal of siRNA by Western blot.

### Resazurin cytotoxicity assay

Mitochondrial activity was assessed by resazurin labeling (Czekanska, 2011). 4T1 and MMT spheroids embedded in 3D collagen were incubated with resazurin-containing media (100 μg/mL), for 4h (at 37°C, 5% CO_2_ humidified atmosphere). Gels were gently washed with PBS and cell-bound fluorescence was measured (ex/em 560/590 nm) using an EILSA reader.

### Real-time quantitative PCR

Collagen gels containing spheroids were dissolved with Trizol (250 μl/100 μl gel; Invitrogen) followed by homogenization prior to chloroform (Merck) phase separation (Farhat, 2012). After centrifugation for 15 min at 12 x 10^3^ g at 4°C, the upper phase was mixed with 70% v/v ethanol and transferred to an RNeasy MinElute spin column for RNA extraction according to the manufactures instructions (Qiagen’s RNeasy kit). After RNA quality control, reverse transcription (SuperScript^®^ II Reverse Transcriptase, Thermo Fisher Scientific) was performed. Primers for conexins and purinergic receptors (Table 1) were designed using HomoloGene (http://www.ncbi.nlm.nih.gov) and were validated *in silico* using Oligoanalyzer software, and *in vitro* using particular positive controls for each gene. qPCR was performed using CFX96 Real-Time PCR Detection System with C1000 Thermocycler (Bio-Rad) and analyzed using the Bio-Rad CFX Manager software (version 2.0). RNA levels for the genes of interest were normalized to the pooled levels of 3 house-keeping genes: β-actin, GAPDH and YWhaz.

**Table 1:**
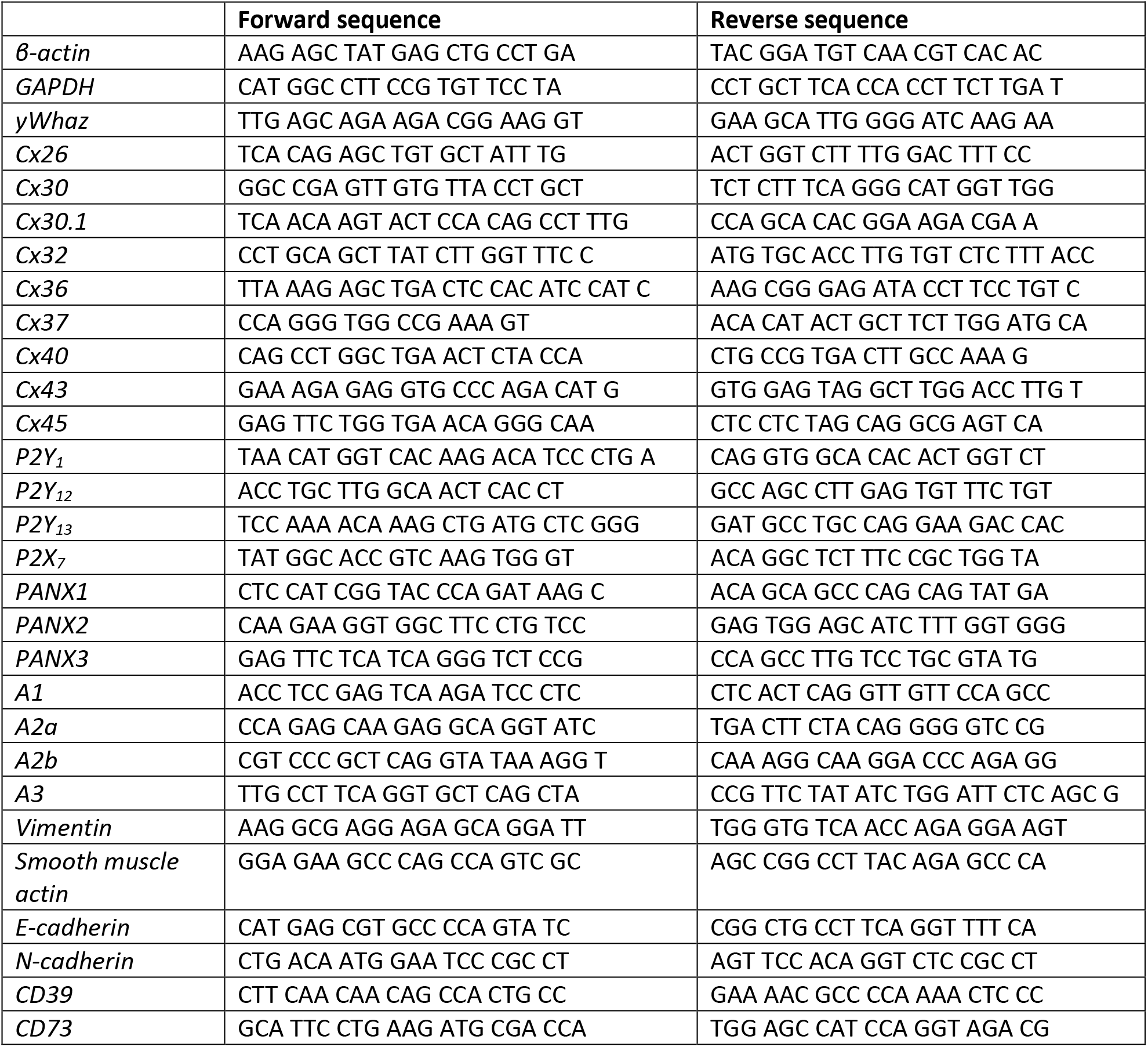
Primer sequences

### Western blot analysis

Cells from 2D culture were lysed using SDS sample buffer (62.5 mM Tris-HCl, 2% SDS, 10% glycerol, 50 mM DTT and bromophenol blue). Collagen gels containing spheroids were digested by collagenase (C0773, Sigma-Aldrich), followed by whole cell lyses using SDS buffer. Western blotting was performed by SDS-PAGE. Briefly, whole cell lysates were loaded and separated on a 10% acrylamide gel with Tris-glycine running buffer followed by blotting to PVDF membrane by wet transfer. PVDF membranes were blotted with monoclonal mouse anti-β-tubulin (E7) (Dep. of Cell Biology, RIMLS, Nijmegen) as loading control; anti-Cx43 (dilution 1:1000); anti-AKT (1:1000); anti-phospho AKT Ser473 (1:500) followed by fluorescence detection (Odyssee; LI-COR Biosciences) and densitometric analysis.

### HPLC analysis for purine content in media

Supernatants from 3D spheroid cultures (24 h) were used for purine analysis. Media contained 80°C heated FCS, which minimizes the degradation of nucleotides and nucleosides (Gendaszewska-Darmach et al., 2003). Adenosine, AMP, ADP and ATP levels were determined by High-Performance Liquid Chromatography (HPLC), similar modified from (Bhatt et al., 2012). In brief, culture supernatant (1 mL) was mixed with chloroacetaldehyde (250 μL; 6x diluted in 1 M acetate buffer, pH 4.5; Sigma-Aldrich), followed by derivatization (60 min, 70°C, 500 rpm) and centrifugation (3 min, RT, 13400 rpm). For HPLC, supernatant (400 μL) was transferred to a HPLC vial and injected. Purines were separated by HPLC (Agilent Technologies 1200 Series) using a Polaris C18-A column (150 x 4.6 mm) with a gradient elution using eluent A (0.1 M K2HPO4, 10 mM TBAHS (pH 6.5), 2% MeOH) and eluent B (H_2_O: ACN : THF; 50:49:1). Retention times were 6.5 (adenosine), 7.7 (AMP), 11.8 (ADP), 15.6 (ATP) and 15.0 min (cAMP). Quantification was based on peak areas of the samples and reference standards were measured with fluorescence detection (excitation: 280 nm; emission: 420 nm).

### Bright-field microscopy

Collective invasion was detected by digital time-lapse bright-field microscopy, as described (Wolf et al., 2013). 3D spheroid cultures were maintained at 37°C and monitored at 5-min intervals for up to 72 h. For pharmacological intervention, spheroids were allowed to migrate for 24 h, followed by addition of inhibitor or vehicle (solvent) for 24 h and wash-out using 1mL of growth media and follow-up for additional 24 h. The number, length and elongation speed of invading strands were quantified using ImageJ (ImageJ; 1.40v; National Institutes of Health). For endpoint experiments, spheroid cultures in 3D collagen after 15, 24 and 48 h of incubation were recorded and the number of pointed and protrusive tips (leader cells) per spheroid was counted manually, as well as length of invading strands were quantified using the image analysis software ImageJ.

### Parachute assay and FACS analysis

Receiver cells (150 x 10^3^ cells/well) were incubated for 24 h (37°C, 5% CO_2_ humidified atmosphere) in 48-well plate to reach 95-100% confluency. Donor cells were loaded with calcein-AM (1 μM, Life Technologies) for 30 min, washed and incubated for further 45 min. (Abbaci et al., 2008). Calcein-labeled cells were detached with EDTA (1 mM) and trypsin (0.075%), centrifuged (251 g, 5 min), re-suspended in media and 75 x 10^3^ cells /well were added over the receiver cell culture at a ratio of 1:2 (donor: receiver) for co-cultured (3, 6 or 12 h). For analysis of dye transfer, cells were detached with EDTA (1 mM) and trypsin (0.075%), assayed by flow cytometry (FACS Caliber) and analyzed using FCS Express software.

### Immunofluorescence staining and confocal microscopy

3D spheroid samples were fixed with 4% paraformaldehyde at room temperature (RT) for 15 min. For detection of intracellular antigens, gels were washed with PBS and permeabilized using 5-10% normal goat serum, 0.3 % Triton-X 100 or 0.2% Saponin in PBS (1h, RT). This was followed by incubation with antibody in PBScontaining 0.1% BSA, 0.3% Triton-X100 or 0.2% Saponin. Antibody dilutions were as follows: Cx43 (1:50), AKT (1:50), S473 pAKT (1:50), β-Catenin (1:200), ZO-1 (1:100), Vimentin (1:200). Primary antibody was incubated at 4°C with shaking overnight, followed by at least 5 washing steps of 15 min each with PBS. Samples were then incubated with secondary antibody (1:400) in PBS buffer for intracellular staining, together with DAPI (5 ug/mL) and phalloidin (1:200) (4 h, RT). Samples were washed at least 5 times (15 min each) and imaged by confocal microscopy (Olympus FV1000; long working distance objectives 20×/NA 0.50, 40×/NA 0.80 or 60x/NA 1.35). Subcellular localization and quantification of Cx43, β-catenin, AKT and pAKT at cell-cell contact and cytoplasmic extensions were determined by manually outlining F-actin positive regions followed by background correction using using ImageJ. For time-lapse confocal microscopy, 3D spheroid cultures consisting of MMT dual color cells or 4T1/H2B-mCherry within 3D collagen were used. Time-lapse microscopy (20x/NA 0.5 air objective LSM 510; Carl Zeiss) with time intervals of 10-15 min for 24 h with usually 4-7 z-scans comprising all of the strands per spheroid (approx. 100 μm in depth). The confocal slices were reconstructed as maximum intensity projections and image analysis including leader cell tracking were performed using ImageJ.

### Gap Fluorescence recovery after photobleaching (GapFRAP)

Cancer cells grown as monolayers on glass bottom dishes or as multicellular spheroids in 3D collagen were labelled with calcein-AM (2-3 μM), in modification from (Kuzma-Kuzniarska et al., 2014). Regions of interest (ROI) were defined for single cells or small regions and bleaching was performed using 100 % Argon laser output 1mW with 200 iterations using a large pinhole (~8 airy units) and 20 s interval at 37°C, 5% CO_2_ humidified atmosphere over 12 min of follow-up. Calcein transfer between cells seeded on 2D cover glass or within collagen gels was monitored using confocal microscopy (LSM 510; 20x/NA 0.5 air; Carl Zeiss; 37°C, 5% CO_2_). Inhibition of GJIC was achieved using CBX (100 μM) at least 45 min before GapFRAP imaging. Time-resolved bleaching and post-bleaching recovery was quantified using ImageJ as the mean fluorescence intensity. For normalizing fluorescence after photobleaching, cells in non-bleached control regions were coregistered and used for background correction. Detached single cells were used as an internal negative control for GJIC-dependent fluorescence recovery.

### Breast cancer tissue sections immunofluorescence

For multispectral imaging of Cx43 and ADORA1, tissue sections with positive invasion margin were selected from 19 breast cancer patients (Table 2). The use of coded tumor tissue was approved by the institutional review board, according to national law (Nagelkerke et al., 2011). Formalin-fixed paraffin-embedded breast tissue sections (5 μm) were deparaffinized, followed by antigen retrieval using Tris-EDTA buffer for 15 min (95-100°C) and washing step with PBS. Sections were blocked with 5% NGS in 0.2% tween for 1 h at RT. Primary antibody was incubated at 4°C with shaking overnight using the following dilutions: Cx43 (1:100), vimentin (1:200), pan-cytokeratin (1: 200), ADORA1 (1:200).

**Table 2:**
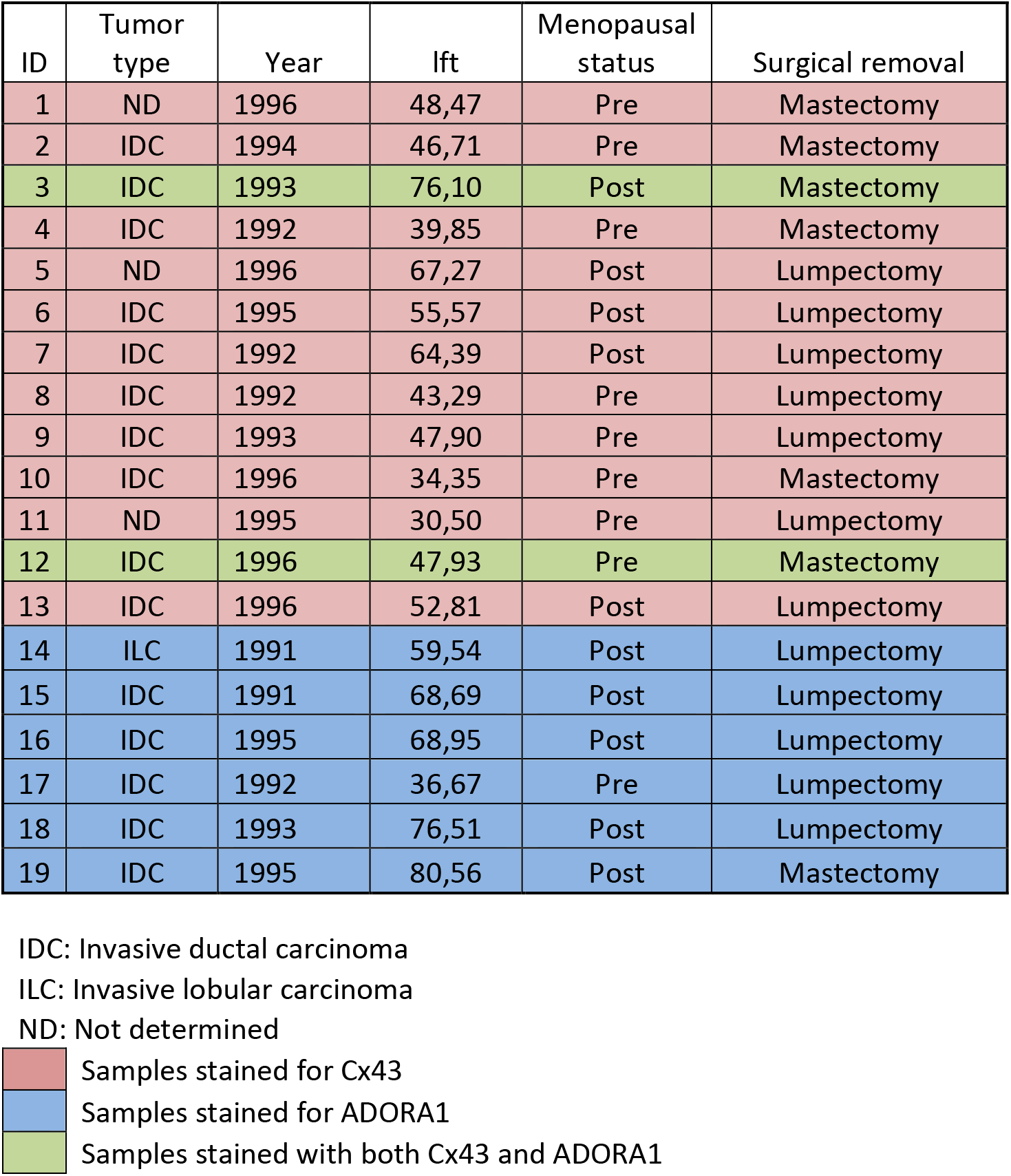
Patients whose tissue sections were stained for Cx43 and/or ADORA1

Tissues were then washed four times (PBS, 5min each), incubated with secondary Alexa-fluor-488/546/647-conjugated goat anti-mouse, -rabbit, and -chicken antibodies, together with DAPI (1 ug/mL), for 1h at RT and further washed four times (PBS, 5min each). Dried sections were embedded in Mowiol (Sigma-Aldrich) and scanned using the automated Vectra Intelligent Slide Analysis System (Version 2.0.8, PerkinElmer).

### Multispectral imaging and quantitative digital analysis

Tissue slides were imaged using Vectra Intelligent Slide Analysis System (Version 2.0.8, PerkinElmer), as described (Mascaux et al., 2019). This imaging technique combines imaging with spectroscopy by collecting the spectral range from 400 to 720 nm in 10 nm steps in an automatic manner. To define the spectrum of each fluorophore as well as of tissue autofluorescence, stained single-fluorescence and unstained tissue was recorded with the Nuance Multispectral Imaging System (Version 3.0.2, PerkinElmer). A spectral library was built from the extracted spectra to enable the quantitative separation and spectral unmixing, thus removing cross-talk between fluorophores and interfering tissue autofluorescence. A selection of 10 representative original multispectral images was loaded into an advanced user-trainable morphologic image analysis software (InForm Version 2.1, PerkinElmer) which utilizes machine learning for user-trained assignment of morphologic and spectral intensity patterns to discriminate tumor cells, stromal regions and normal ducts. The analysis software also utilizes the multimarker, multispectral data to extract and calculate single-cell parameters, including nuclear, cytoplasmic, and membrane intensity values used to classify each cell (e.g. as luminal, cancerous or stroma cell) (Suppl. Fig. 2b). After training, the algorithm was applied as batch analysis of multiple original multispectral images of different samples of the same origin equally stained. Cx43, pan-cytokeratin and vimentin expression levels were quantified in normal bilayered ducts, in tumor cells invading within the fibrous and adipose tissue and in their surrounding stroma. The expression levels of Cx43 were measured in each cell and the positivity threshold was determined by ROC analysis whereby Cx43 values of all the LE cells available in different patients were used as negative control. With 99% specificity and 50% sensitivity, the minimum Cx43 threshold was calculated to be 0.875 (Fig. 4C, dashed line).

### *In silico* analysis of Cx43 and ADORA1 expression in breast cancers

We used the publicly available database (http://kmplot.com), which contains expression data on several genes including Cx43 and ADORA1 in a cohort (580 patients) of basal-subtype breast cancer patients. The database classifies patients, based on the expression levels of selected gene of interest, into high or low expression, to extract correlations between gene expression level and patient outcome.

### Mice

Balb/C female mice (6-8 weeks old) were purchased from the Charles River Laboratories. All animal procedures were approved by the Ethical Committee on Animal Experiments at Radboud University, Nijmegen (protocol number: 140201) and performed according to guidelines of Animal Welfare Committee of the Royal Netherlands Academy of Arts and Sciences, the Netherlands.

### Implantation of tumor cell spheroids in the mammary fat pad

All surgical procedures were performed under isoflurane inhalation anaesthesia (1-2% isoflurane/O_2_ mixture) and performed under conditions of microsurgery, as described (Ilina et al., 2018). In brief, 4T1 dual color spheroids (500 cells/spheroid) were inplantated adjacent to the third mammary fat pad into a central area of the mammary imaging window by nontraumatic microimplantation. After implantation, a sterile cover-glass was placed into the window frame and the integrity of both implanted tumoroid and stroma were verified by fluorescence microscopy.

### Intravital microscopy and image analysis

Tumor growth and cell invasion were monitored for up to 6 days by intravital epifluorescence and multiphoton microscopy, as described (Ilina et al., 2018). In brief, mice were anesthetized with isoflurane and the imaging window was stably fixed on a temperature-controlled stage (37 °C) using a custom holder. Blood vessels were visualized by i.v. injection of 70 kDa dextran labeled with Alexa Fluor 750. Microscopy was performed using multiphoton microscope (LaVision BioTec). Images were analyzed to quantify both, the number of nuclei and the surface/invasion area of implanted tumors at different time points using ImageJ.

### Spontaneous metastasis analysis

4T1 cells (1×10^5^ in 50 μL) were injected into the 4^th^ mammary fat pad of Balb/C 8-week old female mice (Charles River Laboratories) as previously described (Ilina et al., 2018). DMSO and PSB36 (30 mg/Kg) dissolved in PBS (Bilkei-Gorzo et al., 2008) were injected intraperitoneally once per day. Tumor size was recorded weekly by a caliper and mice were euthanized 30-36 days after cancer cell injection or at the humane endpoint (tumor volume of 2 cm^3^). Lungs were harvested, fixed in buffered formalin, embedded in paraffin and sectioned (5 μm-thick). Immunohistochemistry of cytokeratin-8 was used for scoring metastasis and quantification was performed as described (Ilina et al., 2018). The extent of metastasis was quantified as the number of neoplastic events from sequential sections and represented as the number of metastatic foci per slice. Metastasis was classified as micrometastasis, for 3 – 15 cytokeratin-8-positive cells and macrometastasis when more than 15 tumor cells were present in nodular topology.

## Acknowledgements

We gratefully acknowledge Huib Croes, and Dagmar Verweij for technical support and Michael Dustin and Pavlo Grytsenko for helpful discussions. This work was supported by a grant of the German *Excellence Initiative* to the Graduate School of Life Sciences, University of Würzburg (to A.K.), the Netherlands Science Organization (NWO-VICI 918.11.626), the European Research Council (617430-DEEPINSIGHT), and the Cancer Genomics Center, The Netherlands (to P.F.).

## Author contributions

A. Khalil and P. Friedl designed the experiments. A. Khalil, O. Ilina, A. Vasaturo, JH. Venhuizen, M. Vullings, V. Venhuizen and A. Bilos performed the experiments. A. Khalil, O. Ilina, A. Vasaturo, JH. Venhuizen, M. Vullings, V. Venhuizen, A. Bilos and P. Friedl analyzed the data. P. Span provided clinical samples. A. Khalil and P. Friedl wrote the manuscript. All authors read and corrected the manuscript.

