## Supplementary figures and movie legends for "Leader cell activity and collective invasion by an autocrine nucleotide loop through connexin-43 hemichannels and ADORA1"

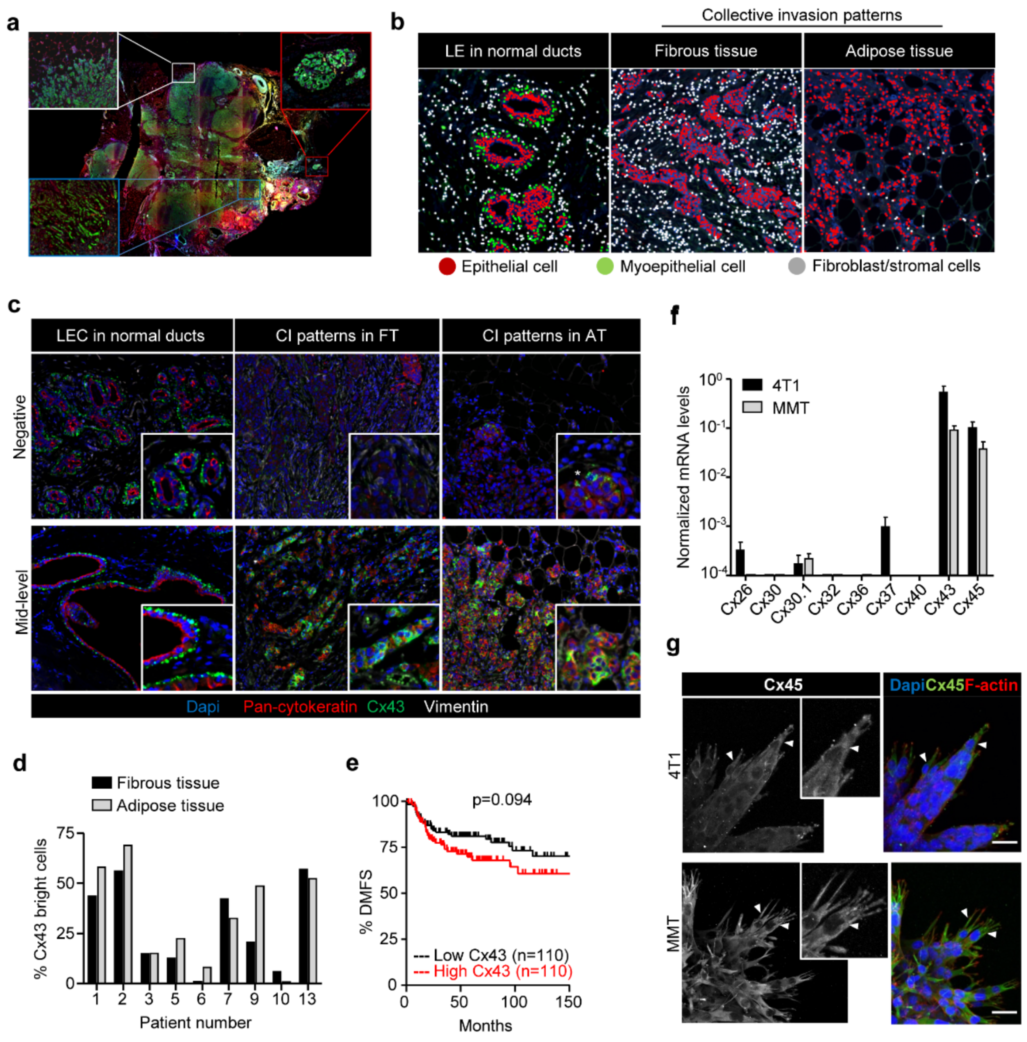


**Suppl. Fig. 1. Expression of Cx43 in breast cancer cells during collective cancer cell invasion.** (a) Original colored fluorescence overview of breast cancer lesions with zoom inserts on normal ducts and tumor invasive margins. (b) Trainable cell phenotyping function of the Inform software to identify different cell types within a multicellular tissue based on spectral fluorescence fingerprinting. Images display the results of the per-cell phenotyping analysis in normal ducts (Luminal epithelial cells, LE), collective invasion (CI) patterns in fibrous and in adipose tissue. In each region, different cell types are marked by colored dots, as indicated. For all regions devoid of bilayered ducts, cells of epithelial origin, based on pan cytokeratin, were considered as invasive breast cancer cells. (c) Unmixed composite images of Cx43 expression in epithelial cancer cells within the tumor margins of two representative patients with negative (upper panel) and mid-level (lower panel) Cx43. (d) Heterogeneous expression of Cx43 within cancer cells of the collective invasion patterns in different tumor regions and samples. Values represent the percentage of Cx43 positive cancer cells. (e) Kaplan-Meier survival plot predicting distant metastasis free survival for high vs low Cx43 expression in basal-type breast cancer patients (Gyorffy B et al.). P values, Log-rank test. (f) qPCR for nine connexin subtypes relevant for the mammary gland (Cx26, Cx30, Cx30.1, Cx32, Cx36, Cx37, Cx40, Cx43 and Cx45). Connexin mRNA expression in 4T1 (black) and MMT (gray) cells from 3D collagen invasion cultures. Values represent mean normalized mRNA levels and SEM from 3 independent experiments. (g) Cx45 localization in collectively invading strands in 3D collagen. Confocal images represent maximum intensity projections from 3D confocal stacks. Bars, 20 μm.

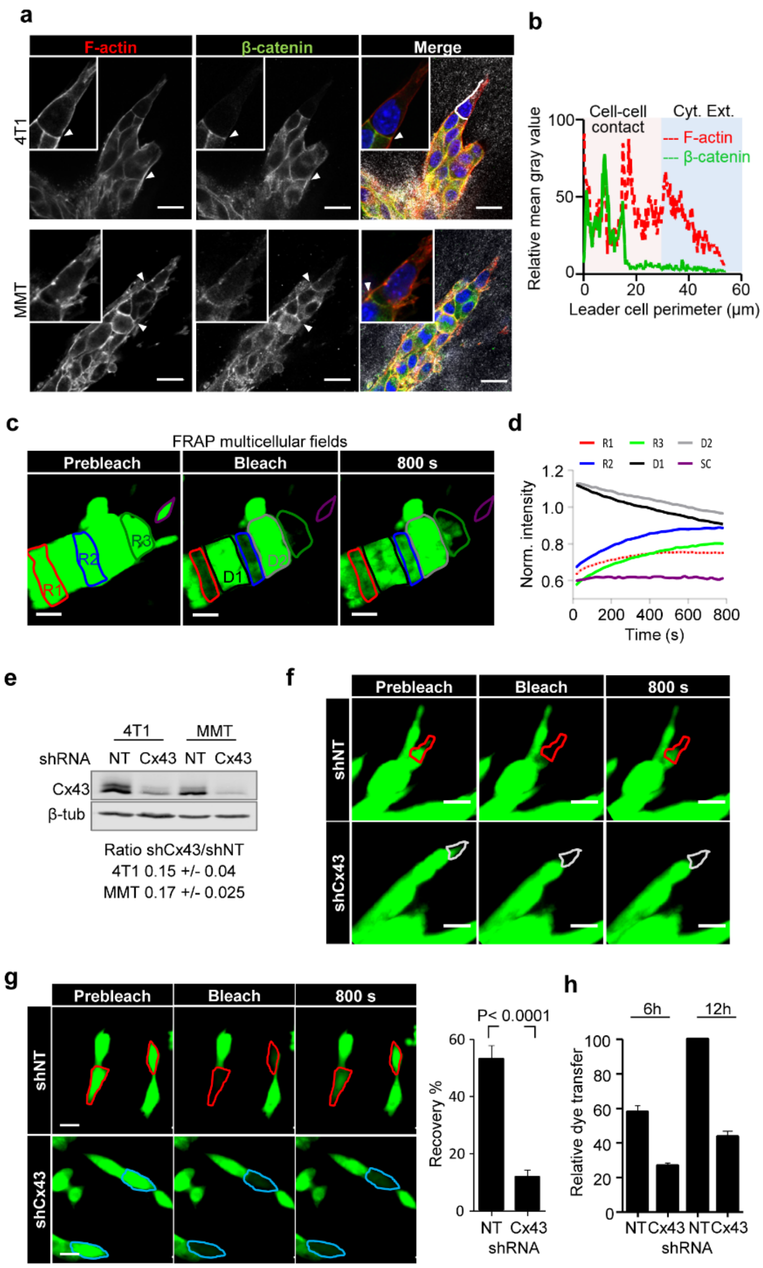


**Suppl. Fig. 2. Collectively invading cancer cells maintain cell-cell connections and communication through Cx43-mediated GJIC.** (a) β-catenin localization along cell-cell junctions between both LC and FC in collective strands of 4T1 and MMT cells invading 3D fibrillar collagen. White arrowheads, cell-cell junctions. (b) Intensity distribution of β-catenin and F-actin along the circumference of a leader cell (white dashed border). (b) Values show the pixel intensity with background subtraction. (c) Multicellular 3D gapFRAP. Single confocal slices of calcein-labeled invasive 4T1 strands before and after photobleaching. Dashed contours represent multicellular regions within an invasive strand. Follower regions (red: R1; blue: R2), invasive front (green: R3), single detached cell (SC: purple), in addition to the unbleached neighboring regions D1 and D2. (d) Normalized mean calcein fluorescence intensity of the selected regions. (e) Western blot for Cx43 and β-tubulin as loading control. Whole cell lysates extracted from 4T1 and MMT cells with stable expression of shNT or shCx43; +/- SD. (f, g) Fluorescence recovery of calcein-labeled MMT leader cells expressing shNT or shCx43 during invasion in 3D collagen (f) and of physically connected cells in 2D culture (g). Values represent the means and SEM of at least 19 bleached cells per condition from 3 independent experiments. P values, Mann Whitney test. (f) Relative dye transfer in parachute assay of 4T1 with non-targeting shRNA (NT) and Cx43 shRNA (Cx43) after 6 or 12 h of cell-cell (donor-receiver) encounter. Values represent the means and SEM from 2 independent experiments. Bars, 20 µm.

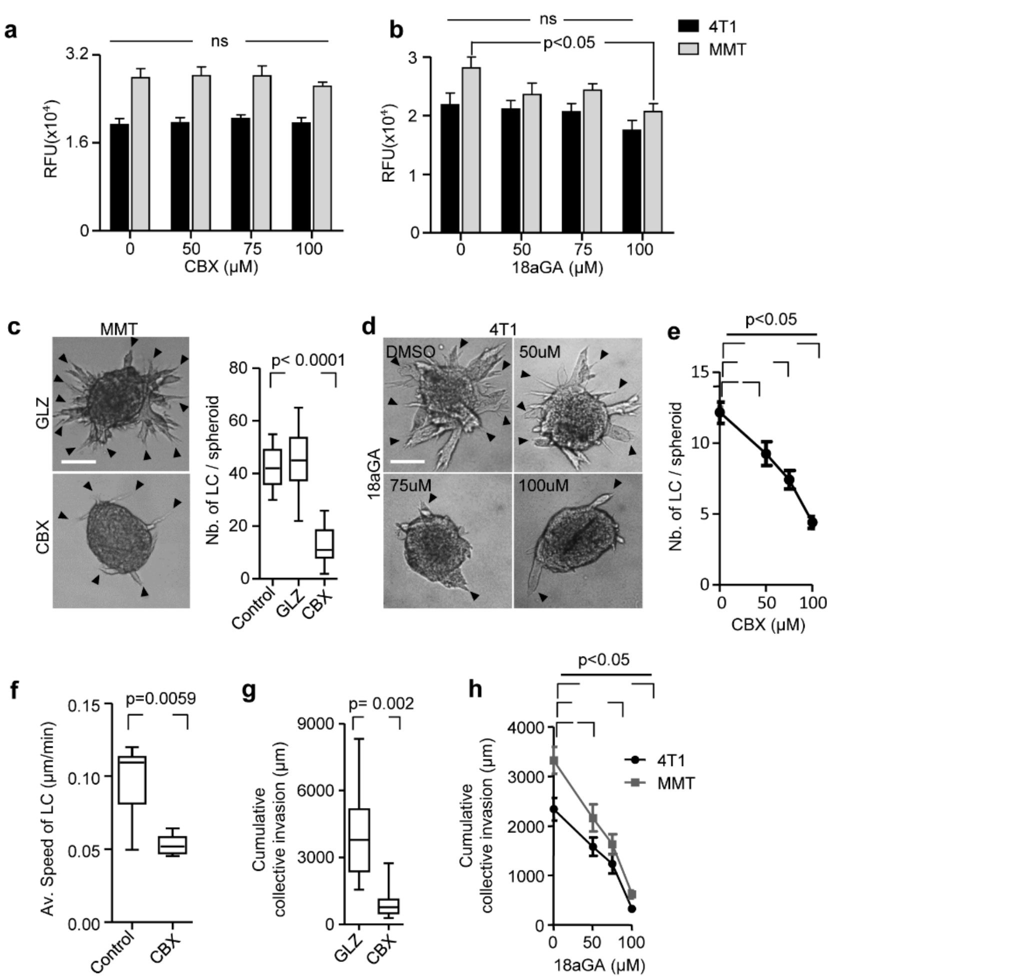


**Suppl. Fig. 3. Cx43 dependence of leader cell function.** (a, b) Resazurin cytotoxicity assay for 4T1 and MMT cells in 3D collagen for 24 h in the presence of vehicle control and escalating doses of CBX (a) and 18aGA (b). Values represent the mean relative fluorescence unit (RFU) with SEM of 2 - 3 independent experiments. P values, one-way ANOVA. (c) Bright-field images after 24 h of invasion in 3D collagen in the presence of CBX or an inactive structurally homologous control compound (GLZ). (c) Leader cell initiation in MMT spheroids after 24 h of culture. (c) Data represent the medians (black line) of 8- 11 MMT spheroids per condition from 3 independent experiments. P value, Mann Whitney test. (d) Efficacy of collective invasion in 4T1 spheroids in the presence of escalating doses of 18aGA. Bright field images after 24 h of culture. (e) 4T1 leader cell initiation in the presence of escalating doses of CBX. Data represent mean values and SEM from 23-40 spheroids per condition from at least 3 independent experiments. P value, ANOVA with Bonferoni multiple comparison test. (f) Speed of MMT leader cells during collective invasion in 3D collagen. Values represent median (black line) of 7-8 LCs tracked. P value, Mann Whitney test. (g) Cumulative collective invasion of MMT cells. Data represent the medians (black line) of 8- 11 MMT spheroids per condition from 3 independent experiments. P value, Mann Whitney test. (h) Cumulative collective invasion of 4T1 and MMT spheroids in the presence of escalating doses of 18aGA. Data represent mean values and SEM from 12 spheroids per condition from 3 independent experiments. P value, ANOVA with Bonferoni multiple comparison test. Bars 50 μm.

**a**

| | Dunnett's Multiple Comparison Test | | | | | | | |
|---|---|---|---|---|---|---|---|---|
| Nucleoside/nucleotide | ADO | | AMP | | ADP | | ATP | |
| Cell line | 4T1 | MMT | 4T1 | MMT | 4T1 | MMT | 4T1 | MMT |
| Ctrl vs CBX | ns | ** | * | ns | ** | *** | ns | ns |
| Ctrl vs Cx43sh | * | ns | ** | * | ** | *** | ns | ns |
| Ctrl vs CBX+Cx43sh | * | ** | ** | ** | ** | *** | ns | * |

* p<0.05 ** p<0.005 *** p<0.0001

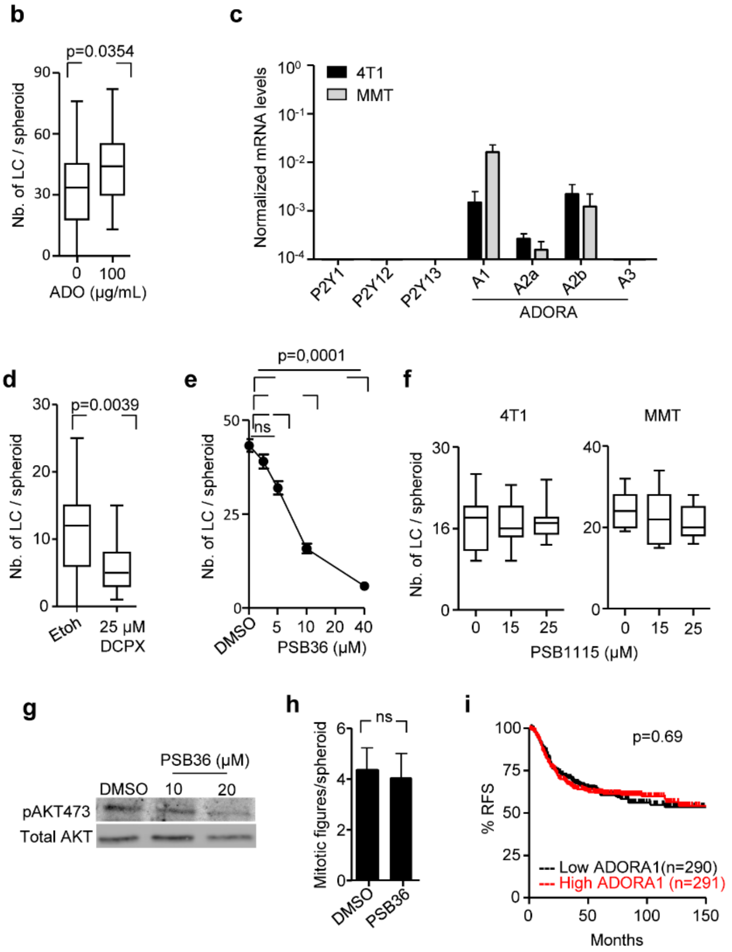


**Suppl. Fig. 4. Cx43-mediated nucleotide release and ADORA in LC initiation. (a)** Summary of the effect of Cx43 downregulation and/or CBX treatment on the reduction of individual nucleotide and nucleosides concentration detected in the media of 4T1 and MMT collagen cultures by HPLC. Concentrations from CBX-treated and Cx43 shRNA cultures were compared to control cells (Ctrl: untreated cells with non-targeting shRNA). P values Anova with Dunnets multiple comparison test. (b) Leader cell initiation in MMT spheroids expressing shCx43 after 24 h of invasion in collagen in the presence and absence of exogenous ADO. Values display median (black line), 25/75 percentiles (boxes) and maximum/minimum values (whiskers) of 30-35 spheroids per condition from 3 independent experiments. P value, Mann Whitney test. (c) Purinergic receptors mRNA expression in 4T1 (black) and MMT (gray) cells during collective invasion in 3D collagen assessed by qPCR. Values represent the mean normalized mRNA levels and SEM from 3 independent experiments. (d) Leader cell initiation in response to DCPX treatment 12-15 h of invasion of 4T1 cells in collagen. Values display medians (black lines) 25/75 percentiles (boxes) and maximum/minimum values (whiskers), 23-27 spheroids from 3 independent experiments. P values, Mann Whitney test. (e) Leader cell initiation in the presence of vehicle and escalating doses of PSB36. Values represent mean values and SEM of at least 20 MMT spheroids per condition from 3 independent experiments. P values, ANOVA with Bonferoni multiple comparison test. (f) Leader cell initiation of 4T1 and MMT spheroids after 12-15 h of invasion in collagen. Values display medians (black lines) 25/75 percentiles (boxes) and maximum/minimum values (whiskers), 4-12 spheroids from one experiment. (g) Western blot for phospho-AKT473, AKT and β-tubulin as loading control. Whole cell lysates extracted from 4T1 spheroids **Suppl. Fig. 4. (continued)** after 24 h of invasion into 3D collagen in the presence of DMSO, 10 and 20 µM of PSB36. (h) Mitotic figures in 4T1 spheroid after 24 h of invasion in collagen. Values represent average number and SEM of mitotic figures per spheroids, 3 spheroids (at least 1000 cell/spheroid) per condition. (i) Kaplan-Meier survival plot predicting relapse free survival (RFS) in basal-type breast cancer patients (Gyorffy B et al.) for high vs low ADORA1 expression. P values, Log-rank test.

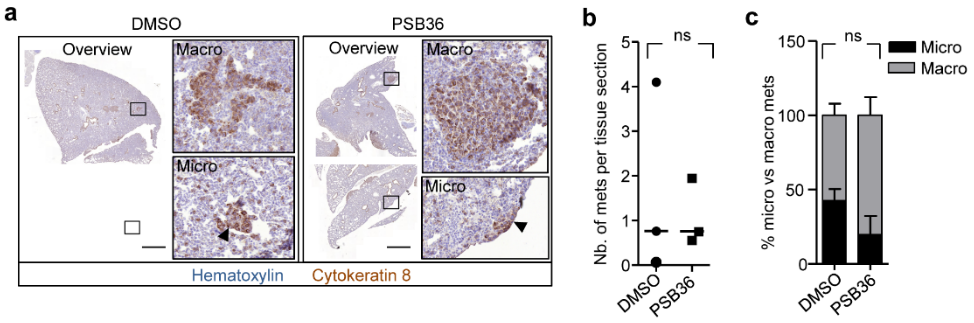


**Suppl. Fig. 5. Effect of ADORA1 antagonist treatment on metastasis to the lungs *in vivo*.** (a) Overviews of lung sections isolated from DMSO and PSB36 (30mg/kg) treated mice, stained for cytokeratin 8. Insets and arrowheads depict macro- and micrometastasis. (b) Cumulative number of micro- and macrometastases normalized to the number of lung sections analyzed. Data represent a total of 135-145 lung sections from 3 independent mice per group. (c) Absolute counts of micro- or macrometastasis per treatment group: DMSO: 39 micro, 59 macro, 98 total; PSB36: 39 micro, 91 macro, 130 total. Data show the means and SD. P value, Mann Whitney test. Bars, 1000 µm.

**Supplementary movies**

**Suppl. Movie 1.** Bright-field time-lapse microscopy of 4T1 spheroids showing Protrusive tips during initiation of collective invasion in 3D collagen I. Field size: 529x301 µm.

**Suppl. Movie 2.** Time lapse imaging by confocal microscopy of MMT spheroids (H2b-GFP and cytosolic dsRed) during initiation and elongation of collective invasion in 3D collagen I. Field size: 1463x1463 µm.

**Suppl. Movie 3.** Time lapse confocal imaging of calcein-labeled 4T1 spheroids before and after photobleaching of cells and multicellular regions within invasive strands in the absence (control media) and presence of the gap junction inhibitor (CBX) in 3D collagen I. Field size: 213x122 µm. Related to Fig. 2b and Supp. Fig. 2c.

**Suppl. Movie 4.** Time lapse confocal imaging of collective invasion of MMT cells (H2b-GFP) in 3D collagen I in the presence and absence of CBX. Field size: 540x550 µm. Related to Supp. Fig. 3f.

**Suppl. Movie 5. Intervention with collective invasion by the gap junction channel blocker (CBX).** Time lapse imaging by bright-field microscopy of established invasive 4T1 strands in 3D collagen I after challenge with connexin inhibitor CBX and followed by drug washout. Field size: 1174x881 µm. Related to Fig. 3l.

**Suppl. Movie 6. Intervention with collective invasion by ADORA1 inhibitor (PSB36).** Time lapse imaging by bright-field microscopy of collective invasion of 4T1 cells in 3D collagen I before and after the addition of PSB36. Field size: 1174x881 µm.
